# Exploring anterior thalamus functional connectivity with cortical regions in prospective memory with ultra-high-field fMRI

**DOI:** 10.1101/2024.02.14.580346

**Authors:** Luke Flanagan, Bruno de Matos Mansur, Christoph Reichert, Anni Richter, Soroosh Golbabaei, Jasmin M. Kizilirmak, Catherine M. Sweeney-Reed

**Affiliations:** Neurocybernetics and Rehabilitation, Dept. of Neurology, Otto von Guericke University, Magdeburg, Germany; Leibniz Institute for Neurobiology, Magdeburg, Germany; German Center for Mental Health (DZPG), partner site Halle-Jena-Magdeburg, Germany; Center for Intervention and Research on adaptive and maladaptive brain Circuits underlying mental health (C-I-R-C), Halle-Jena-Magdeburg, Germany; Dept. of Psychiatry and Psychotherapy, Jena University Hospital, Jena, Germany; Neurodidactics and NeuroLab, Institute of Psychology, University of Hildesheim, Germany; German Center for Neurodegenerative Diseases, Göttingen, Germany; Center for Behavioral Brain Sciences, Otto von Guericke University, Magdeburg, Germany

**Keywords:** prospective memory, anterior nuclei of the thalamus, 7T MRI, functional connectivity, strategic monitoring

## Abstract

Prospective memory, or memory for future intentions, engages particular cortical regions. Lesion studies also implicate the thalamus, with prospective memory deterioration following thalamic stroke. Neuroimaging, anatomical, and lesion studies suggest the ANT (anterior nuclei of the thalamus) in particular are involved in episodic memory, with electrophysiological studies suggesting an active role in selecting neural assemblies underlying particular memory traces. Here we hypothesized that the ANT are engaged in realizing prospectively-encoded intentions, detectable using ultra-high-field strength functional MRI. Using a within-subject design, participants (N = 14; age 20-35 years) performed an ongoing n-back working memory task with two cognitive loads, each with and without a prospective memory component, during 7-Tesla functional MRI. Seed-to-voxel whole brain functional connectivity analyses were performed to establish whether including a prospective memory component in an ongoing task results in greater connectivity between ANT and cortical regions engaged in prospective memory. Repeated measures ANOVAs were applied to behavioral and connectivity measures, with the factors *Task Type* (with prospective memory or not) and *N-Back* (2-back or 3-back). Response accuracy was greater and reaction times faster without the prospective memory component, and accuracy was higher in the 2-than 3-back condition. *Task Type* had a main effect on connectivity with an ANT seed, with greater ANT–DLPFC (dorsolateral prefrontal cortex) and ANT–STG (superior temporal gyrus) connectivity when including a prospective memory component. Post hoc testing based on a significant interaction showed greater ANT–DLPFC connectivity (p-FWE = 0.007) when prospective memory was included with the low and ANT–STG connectivity (p-FWE = 0.019) with the high cognitive load ongoing task. Direct comparison showed greater functional connectivity between these areas and the ANT than DMNT (dorsomedial nucleus of the thalamus) during prospective remembering. Enhanced ANT–DLPFC connectivity, a brain region with an established role in strategic monitoring for prospective memory cues, arose with a low cognitive load ongoing task that enabled monitoring. This connectivity was significantly less on direct comparison with increasing the cognitive load of the ongoing task without prospective memory, suggesting specificity for prospective memory. Greater ANT–STG connectivity on prospective memory inclusion in the higher cognitive load ongoing task fits with reported STG activation on prospective memory through spontaneous retrieval. Lower connectivity on direct comparison with a DMNT seed suggests ANT specificity. The findings fit with a coordinating role for the ANT in prospective remembering. Given the small sample, these findings should be considered preliminary, with replication required.

## Introduction

Prospective memory, the ability to remember and fulfill future intentions^1–3^, plays a pivotal role in essential tasks such as adhering to medication schedules and attending appointments. Notably, a decline in prospective memory constitutes one of the most commonly reported memory concerns^4^. Prospective memory undergoes impairment with advancing age and is prevalent in conditions such as mild cognitive impairment and Parkinson’s disease^5–8^, underlining the importance of understanding the neural processes involved in a young, healthy population for the future development of effective diagnostic and therapeutic strategies when prospective remembering is impaired, which will involve targeting the relevant deficits in a given patient group.

Prospective remembering involves a complex integration of multiple systems, including processes underlying episodic and working memory and also attentional processes^9–11^. Prospective memory can be time-based, when a specific action is to be performed at a particular future time point, or event-based, when a prospectively encoded intention should be carried out on detection of a specified cue^5,12,13^. Here we focused on the latter, in which the timing of prospective memory events and their neural correlates can be more precisely controlled. Investigation of event-related prospective memory classically involves the intention to perform a particular action, after a delay, on detection of a prospectively-encoded prospective memory cue^5^. An important aspect is the performance of an ongoing task that prevents continuous rehearsal of the prospective memory task^5,14^. Based on engagement of different brain networks supporting key aspects of a prospective memory task, neurocognitive models have been proposed to explain how prospectively-encoded future intentions are realized. An influential model of how prospective memory is achieved is by either strategic monitoring of the environment for the prospective memory cue, which involves a negative impact on ongoing task performance measures, by spontaneous retrieval of the prospective memory task on prospective memory cue detection, or by a combination of these approaches^15^. The extent to which a particular approach is favored depends on the cognitive load involved in the ongoing task and prospective memory cue features, such as the focality, saliency, and frequency^11,15–17^. When strategic monitoring is employed, the prospective memory component may be maintained in working memory, as the environment is monitored for a prospective memory cue during ongoing task performance^18^. When the prospective memory task is achieved through spontaneous retrieval, the action to be performed in the future is not continuously actively maintained, and the prospective memory is deemed to be stored in long-term, episodic memory rather than being held in working memory^19^.

We used the n-back as the ongoing task, as in previous studies of prospective memory, with infrequent cues in a different color requiring a different response^9,20^. We chose this study design for several reasons. Firstly, the cognitive load can be readily manipulated, enabling both modulation of the approach that participants use to achieve the prospective memory task component and examination of whether potential neural correlates of the prospective memory task component reflect prospective memory or simply an increase in cognitive load. While the performance of an ongoing task and a prospective memory task may be considered to be a dual task paradigm^18^, several criteria have been proposed to distinguish between a prospective memory and a classical dual task paradigm, in which two tasks are actively maintained and performed in parallel: 1) the cognitive load of the ongoing task in a prospective memory paradigm should be sufficient to prevent continuous active rehearsal of the prospective memory component^5,14,21^; 2) a prospective memory paradigm requires a delay between the formation of the prospective intention and its performance^12^; 3) prospective memory items should appear less frequently than ongoing task items^16^; 4) prospective memory testing requires task switching, with inhibition of the ongoing task response^22,23^. A rate of 10% of the trials being prospective memory has been proposed in previous studies of event-related prospective memory, including when an n-back task is used as the ongoing task^16,21^⊡. A further reason for choosing this study design is that it includes multiple prospective memory trials, rendering adequate trial numbers for functional connectivity analyses. To avoid appearance of prospective memory cues serving as a repeated reminder of the prospective memory task, no feedback was provided^9^.

Neuroimaging and electrophysiological investigations consistently highlight the involvement of frontal and parietal cortical regions in prospective remembering^10,11,24^. Specifically, the dorsolateral prefrontal cortex (DLPFC) is engaged in both the maintenance and retrieval phases of prospective memory tasks^25–27^. Activation of a dorsal frontoparietal network is consistently observed during prospective memory maintenance and deemed to reflect top-down attentional processes towards external prospective memory cues and the content of prospectively-encoded intentions^10,11^. Neuroimaging meta-analysis has also shown engagement of the thalamus in prospective remembering^10^, and prospective memory deterioration has been reported following thalamic stroke^28^.

Neuroimaging and anatomical studies suggest the anterior nuclei of the thalamus (ANT) in particular are involved in episodic memory processing^29–32^. The ANT have been proposed to be part of an extended hippocampal system, based on their reciprocal anatomical connectivity with the hippocampus and their firing in the theta frequency range, the dominant hippocampal rhythm^29,33^. Lesion studies also indicate ANT involvement in episodic memory^28,34,35^. Evidence from intracranial recordings from the ANT of epilepsy patients suggests that the ANT play an active role in selection of neural assemblies underlying particular memory traces^31,32,36–38^. Prospective remembering requires integration of attentional processes with activity in the brain networks underpinning episodic memory for prospective memory cues and prospectively encoded intentions. Based on the extensive anatomical connectivity between the ANT and widespread cortical regions^29,36^, we postulated that the ANT are engaged in coordinating the cortical activity in attentional and episodic memory networks, to reactivate a prospectively encoded intention on prospective memory cue detection.

Here we performed a seed-based functional connectivity analysis using a left ANT seed in a seed-to-voxel whole brain analysis to examine whether functional connectivity with cortical regions known to be engaged in the attentional processes underlying prospective remembering is greater during performance of a task with a prospective memory component than during a task condition comprising the ongoing task alone. We hypothesized that functional connectivity with the ANT would be detectable based on blood oxygen-dependent (BOLD) measurements acquired using 7-Tesla (7T) functional magnetic resonance imaging (fMRI) employing an event-related design. Ultra-high magnetic field strength scanning results in a better signal-to-noise ratio than using a low strength magnetic field, with a supralinear relationship between magnetic field strength and signal-to-noise ratio^39^. Comparing using 7T with 3T field strength, subcortical structures in particular have been shown to be better discernable at the higher field strength^40^. Moreover, a 7T field strength has enabled detection of memory traces in the medial temporal lobe as well as differentiation between hippocampal subfields during associative learning^41,42^. The left ANT seed location was derived from a previous fMRI study investigating the role of the ANT and also the dorsomedial nucleus of the thalamus (DMNT) in episodic memory processing^30^. The hypothesis was based on a left ANT seed, because our paradigm, like that used by Pergola et al. (2013), can be performed using verbal-based strategies, and verbal processing is well-established as being lateralized to the left hemisphere^43^. Moreover, left lateralization has been associated with prospective memory processing on neuroimaging meta-analysis^27^. To assess the relevance of laterality, the seed location was mirrored to the right side for comparison. Functional connectivity with a left DMNT seed was also examined, to evaluate the specificity of the engagement of the ANT. The classical n-back working memory task was used as the ongoing task^44–46^, because the cognitive load can be readily adjusted, enabling assessment of whether differences in functional connectivity arising on adding the prospective memory task result from increased cognitive load rather than prospective memory processing.

## Materials & Methods

### Participants

We were not aware of a directly comparable 7T fMRI study from which to conduct a power analysis to determine a suitable participant sample size. However, we identified a resting state 7T fMRI study in which seed-based whole brain functional connectivity, using seeds in subthalamic regions, was applied to compare connectivity in two participant groups^47^. We used G*Power to calculate the sample size based on this study^48^. Based on the effect sizes of 1.90 and 1.85 provided for a connectivity difference between a subthalamic region and prefrontal cortex (the most anteriorly listed region) on the left and right respectively, which we selected based on evidence for prefrontal engagement in prospective memory^25–27^, 13 participants would be required to detect a difference at a power of 0.95 with an alpha threshold of p = 0.001 using a paired T-test. An additional power calculation based on our within-subject, repeated measures study design with four repeated measurements, with a medium effect size of 0.6, showed that 14 participants would be required to detect a difference at a power of 0.95 with an alpha threshold of p = 0.001.

Fourteen right-handed (self-reported) healthy participants aged 20-35 years were recruited among students at the Otto von Guericke University, Magdeburg. The mean age of the participants was 28.6 years (SD 3.3; 7 female). Participants all had normal or corrected to normal vision and no history of psychological or neurological disorders, regular medication, or recreational drug use. The study was approved by the Local Ethics Committee of the University Hospital, Magdeburg, in accordance with the Declaration of Helsinki. All participants provided informed, written consent prior to inclusion in the study and were informed of their right to cease participation at any time without providing reasons. Standard MRI exclusion criteria were applied.

### Behavioral paradigm

Visual stimuli were presented using Presentation software (Version 23.1 Build 09.20.22, Neurobehavioral Systems, Berkeley, CA, USA) using back-projection. The task comprised the n-back working memory task as the ongoing task, with a color-based prospective memory task. Each trial comprised 500 ms presentation of a letter and a jittered 1800-2100 ms response period. A sequence of letters was presented on a screen, and participants pressed one button, using a button box, when the current letter corresponded with the letter n letters previously and another button if not (Fig. 1A). The buttons were counterbalanced across participants. Each letter was presented in one of four colors (red/blue/green/yellow, specified in the RGB color space), assigned independently of the letter itself. The prospective memory cue was a particular color, in response to which a third button was pressed with the thumb, instead of making an ongoing task (n-back) response. In other words, the prospective memory cue indicated a brief task switch restricted to the respective trial, irrespective of which n-back response would have been correct. The thumb was used for the prospective memory answer in all participants, because counterbalancing would require using a thumb and one finger for a two-button task (the n-back task), which would be unusual and likely serve as a continual reminder of the prospective memory task. Responses in an n-back task typically use the index and middle fingers^9,20,49^. All four cue colors were included in every block, irrespective of whether a color was assigned as a prospective memory cue in a particular block, so the only difference between prospective memory blocks and ongoing task blocks was the assignment of a particular color as the prospective memory cue for a prospective memory block.

**Figure 1.**
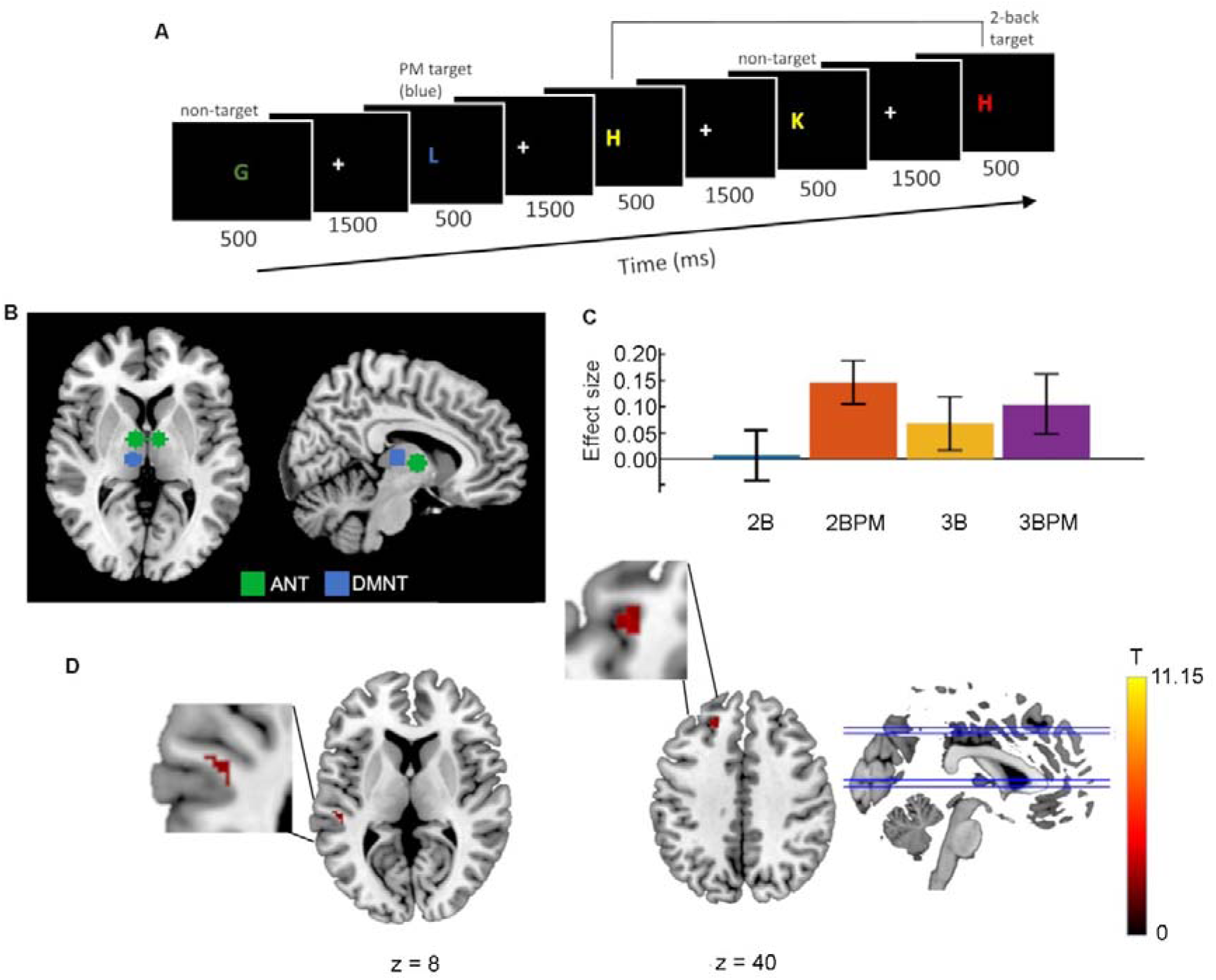
Functional connectivity with the left anterior nuclei of the thalamus during prospective remembering. **A**. Behavioural paradigm. Illustration of the 2-back task, with the colour blue as the prospective memory cue. **B.** Seeds in the left and right anterior nuclei of the thalamus (green) and the left dorsomedial nucleus of the thalamus (blue). **C**. Effect sizes in the four task conditions at the cluster locations where a main effect of *Task type* was observed. 2B: 2-back only; 2BPM: 2-back with prospective memory component; 3B: 3-back only; 3BPM: 3-back with prospective memory component. Cluster MNI coordinates are given below. **D**. Functional connectivity based on a seed in the anterior nuclei of the thalamus (Seed 1), with clusters showing the main effect of *Task Type*. Greater connectivity in the conditions including a prospective memory component can be seen in the left dorsolateral superior frontal gyrus (ANOVA: p-FWE = 0.007) and the left superior temporal gyrus (ANOVA: p-FWE = 0.019). T represents the T-values relating to the cluster size threshold. ANT: anterior nuclei of the thalamus; DMNT: dorsomedial nuclei of the thalamus.

Written instructions were provided to explain how to perform the four task types. The importance of the ongoing n-back task was emphasized. The participants were told that in some blocks they would be shown a particular letter color at the beginning of the block, which would require pressing a different button with the thumb. They were informed that this additional instruction was only valid for performing that block, and the n-back response with the other two buttons would be required for all other trials and blocks. In particular, on starting a new block, a color previously used as a prospective memory cue was no longer relevant, and an n-back response should be given. Participants first performed the four task types outside the scanner to allow an opportunity to ask any questions and to minimize potential training effects in one block type affecting subsequent performance in a subsequent block type, as n-back performance improves with practice, and the improvements are greatest early in task performance^45^. The task comprised four variants: 2-back without a prospective memory cue (2B); 2-back with a prospective memory cue (2BPM); 3-back without a prospective memory cue (3B); 3-back with a prospective memory cue (3BPM). Each variant was performed four times, yielding a total of 16 runs per participant. The runs were carried out in a pseudorandom order, counterbalanced across participants, such that participants could perform low or high cognitive load variants first, as well as variants with or without prospective memory component first. Each run comprised 120 trials (lasting ∼4 min). Based on previous studies, the prospective memory conditions included 10% prospective memory trials^16,21^. The ongoing task target rate of the current study was 15%.

Accuracy was quantified using d’, a sensitivity measure based on the signal detection theory, given by the difference between z-scores of the proportions of hits and false alarms^50^. The accuracy and reaction times (RTs) were analyzed using SPSS (IBM, New York). Mean d’ and RTs were calculated for each participant. Two-way repeated measures ANOVAs were applied, with the within-subject factors *N-back* (2-Back, 3-Back) and *Task type* (with, without prospective memory), followed by post hoc tests. For effect size measures, we computed partial eta squared (η_p_^2^). We also applied two-sided paired T-tests to compare the percentage of prospective memory items with a correct prospective memory response and the RTs to these items in the 2- and the 3-back conditions and to compare RTs to ongoing task and prospective memory items.

To evaluate whether the 3-back was perceived to be more challenging than the 2-back condition, and whether adding a prospective memory task further increased the challenge, participants were asked to rate their ability to perform each of the four block types, using scales from 1 reflecting “mostly guessing” to 5 reflecting “no mistakes”. The reported guessing rate provided a proxy to establish the perceived difficulty of the task, reflecting the likely cognitive load as well as the reliance on spontaneous retrieval rather than strategic monitoring for the prospective memory task component. We interpreted a higher reported rate of guessing as reflecting a greater reliance on spontaneous retrieval. Participants were also asked to report their approach to the prospective memory task component on a scale from 1, indicating spontaneous retrieval, to 5, indicating strategic monitoring. The scale was provided at the end of the whole experiment, as presenting it after each condition type would have emphasized the prospective memory task, potentially precluding subsequent spontaneous retrieval. For the latter scale, answers indicating that the participants were mostly or predominantly using spontaneous retrieval for the prospective memory task component were pooled as spontaneous retrieval (n = 3), answers in the middle of the scale were considered to reflect equal application of the two approaches (n = 4), and answers indicating a mostly or predominantly monitoring approach were pooled as strategic monitoring (n = 3). The scales were introduced after the start of the study, following discussions with early participants, resulting in twelve participants responding to the approach scale and ten to the ability scales. In addition to visualization, paired T-tests were applied to compare participants’ perceived ability according to cognitive load in the ongoing task and to whether a prospective memory task component was included, based on simulation studies evaluating application of parametric statistics to Likert scale data^51,52^.

To examine whether self-reported predominant approach to the prospective memory task had an impact on performance, we applied two-way repeated measures ANOVAs to d’ and to RT with the within-subject factors *Task Type* (with, without prospective memory) and *N-Back* (2-back, 3-back), including the covariate *Reported Approach*. We also applied one-way repeated measures ANOVAs to the hit rates and RTs to prospective memory cues, with the within-subject factor *N-Back*, correcting for the covariate *Reported Approach*.

Finally, to assess whether a training effect resulted in overall improved n-back performance over time, we applied two-sided paired T-tests to compare the d’ and RT in the first and last runs of the blocks, averaging across block type. So that training during one block type would not lead to better performance in a subsequently performed block type, we counterbalanced the order of the block types. To evaluate whether this approach was effective, we applied two-way repeated measures ANOVAs to the changes in d’ and RT over the course of the blocks with the within-subject factors *Task Type* and *N-Back*.

### fMRI data acquisition

Structural and functional MRI data were recorded using a 7T MR scanner (Siemens MAGNETOM 7TPlus at the MRI Core Facility of the Otto von Guericke University Magdeburg). The head coil (Nova Medical) had 32 receiving channels and one transmission channel. Shimming was applied over the whole head using B0 shim mode: Brain and B1 Shim Mode: Trueform. First, a T1-weighted magnetization-prepared rapid gradient echo (MPRAGE) image (TR = 2300 ms; TE = 2.73 ms; flip angle = 5°; 224 sagittal slices; FOV: 256 x 256 mm²; voxel size = 0.8 x 0.8 x 0.8 mm^3^) was acquired. This MPRAGE was used for the two-stage registration of the functional scans to the Montreal Neurological Institute (MNI) space. During the fMRI sessions, T2*-weighted echo-planar images (EPI; TR = 2000 ms; TE = 25 ms; flip angle = 75°; 81 axial slices; FOV: 212 x 212 mm²; voxel size = 1.5 x 1.5 x 1.5 mm^3^) were acquired, employing 3x multiband acceleration with GRAPPA parallel imaging (acceleration factor 2). Slice acquisition was interleaved, in an ascending order, and the phase encoding direction was anterior to posterior. Fat suppression was not applied. During each of the sixteen 230 s runs, 115 volumes were acquired. All scans featured EPI distortion correction^53^. The first six scans during fMRI start-up were discarded. Participants were visually monitored to exclude severe motion during scanning. The mean total scanning time per participant was 1:25:12 h.

### Functional connectivity analysis

Analyses were performed using the Functional Connectivity Toolbox (CONN: Version 22^54,55^ and SPM-12: Statistical Parametric Mapping 12)^56^, using a Windows operating system.

#### Seed creation

The seeds, with 5-voxel radii (1 voxel = 1.8 mm), were created for each seed location in FSLeyes^57^ (Fig. 1B). The location identified by Pergola *et al.* (2013; MNI: −3 −6 3) is located by the anterolateral border of the thalamus with the ventricle. As the volume of any seed at these coordinates, including a single voxel radius, created in FSLeyes using the MNI 2 mm brain, would overlap with the ventricle, the seed was placed five voxels medially, such that it was located entirely within the anterior thalamus. Seed 1 (MNI: −8 −6 3) is localized to the left anterior thalamus (Automated Anatomical Labelling atlas, version 3: AAL3^58^, and the anterior–posterior location did not differ from that reported by Pergola et al. (2013) for the ANT. The seed location was also mirrored to the right side to examine potential laterality. While two other sets of ANT coordinates associated with memory retrieval were given, these locations were not localized by the AAL3 atlas to the ANT. Finally, a seed location in the DMNT from the same study (MNI: −6 −19.5 7.5), confirmed by AAL3 to be in left mediodorsal thalamus, was also used, to enable examination of whether any functional connectivity identified with Seed 1 was specific to the ANT.

#### Pre-processing

Functional and anatomical data were pre-processed using a standard pre-processing pipeline, including realignment with correction of susceptibility distortion interactions, slice time correction, outlier detection, segmentation and normalization to the MNI space, using 4^th^ order spline interpolation to re-slice functional data to yield 1.8 mm isotropic voxels, and smoothing^59^. Slice position was landmark-based, according to the AC–PC line. Functional data were realigned using the SPM realign and unwarp procedure, where all scans were co-registered to a reference image (first scan of the first session) using a least squares approach and a 6-parameter rigid body transformation, and resampled using b-spline interpolation to correct for motion and magnetic susceptibility interactions^60,61^. Potential outlier scans were identified using Artifact Detection Tools, as acquisitions with framewise displacement above 0.9 mm or global BOLD signal changes above 5 standard deviations^62–64^, and a reference BOLD image was computed for each participant by averaging all scans excluding outliers. Functional and anatomical data were normalized into standard MNI space and segmented into grey matter, white matter, and cerebrospinal fluid tissue classes using the SPM unified normalization and segmentation algorithm^65,66^ and the IXI-549 tissue probability map template provided with CONN. Finally, the functional data were smoothed using spatial convolution with a Gaussian kernel of 3 mm full-width half maximum.

The functional data were denoised using a standard denoising pipeline. The steps included the regression of potential confounding effects characterized by white matter time series (5 CompCor noise components), cerebrospinal fluid time series (5 CompCor noise components), motion parameters and their first order derivatives (12 factors^59,67^), outlier scans (below 14 factors^63^), session and task effects and their first order derivatives (8 factors), and linear trends (2 factors) within each functional run. Considering prior studies highlighting the confounding influence of main task effects on functional connectivity outcomes^68,69^, the main task effect was modeled as a response convolved with the canonical hemodynamic response function, along with its first-order derivative. Both were subsequently regressed out to mitigate task-related modulation and isolate functional connectivity results. Regression was followed by bandpass frequency filtering of the BOLD time series^70^ between 0.009 Hz and 0.08 Hz, as applied previously in fMRI studies using the n-back task^71,72^. CompCor^73,74^ noise components within white matter and cerebrospinal fluid were estimated by computing the average BOLD signal as well as the four largest principal components orthogonal to the BOLD average, as recommended based on an investigation of noise reduction including different numbers of principal components and evaluating anticorrelations between resting and task-based networks^74^. From the number of noise terms included in this denoising strategy, the effective degrees of freedom of the BOLD signal after denoising were estimated to range from 143.7 to 438.9 (average 397.8) across all participants^64^.

#### First level analyses

First-level analysis seed-based connectivity maps were estimated to characterize the spatial patterns of functional connectivity associated with a predefined seed region. Using the CONN toolbox, weighted seed-based connectivity maps were computed by correlating the average BOLD time series of the seed region with the time series of all other voxels in the brain for each task condition. This approach enables characterization of condition-specific functional connectivity strength. Three seed regions were analyzed (see Seed creation section). Functional connectivity strength was thus represented by Fisher-transformed bivariate correlation coefficients from a weighted least squares general linear model^59^, estimated separately for each seed area and target voxel, modelling the association between their BOLD signal time series. Individual time series were weighted by a boxcar signal characterizing each task or experimental condition convolved with an SPM canonical hemodynamic response function. The weights were defined to encompass entire runs for each condition.

To investigate further whether the observed functional connectivity differences with the ANT when including a prospective memory component, compared with ongoing task performance alone, reflected the inclusion of the prospective memory task rather than a general cognitive load increase, we also compared functional connectivity when including the trial onsets as a regressor. While the neural correlates of cognitive processing accompanying strategic monitoring are sustained throughout a prospective memory paradigm, the correlates of spontaneous retrieval are transient, arising when a prospective memory cue is detected^10,24^. A psychophysiological interaction (PPI) analysis was conducted to evaluate the effect of prospective memory on the seed to voxel connectivity of the left ANT. PPI analysis is an approach to the investigation of task-specific changes in the relationship between activity in different brain areas, such as a seed region and other voxels, for example, as here, in the whole brain, in a particular psychological paradigm^75^. To this end, we took into account the prospective memory item onset times. This analysis enabled evaluation of whether functional connectivity observed when a prospective memory component was added was temporally associated with the appearance of prospective memory cues.

#### Second level analyses

Group-level analyses were performed using a general linear model framework, specifically employing a linear mixed-effects model to account for both fixed effects of the experimental conditions and random effects across participants. This approach allows for the analysis of voxel-level hypotheses by incorporating multivariate parametric statistics, which include variability across participants as random effects. Additionally, sample covariance estimation was used to account for variability across multiple outcome measures. Inferences were performed at the cluster level, based on parametric statistics from Gaussian Random Field theory^59,76^. Results were thresholded using a combination of a cluster-forming p < 0.001 voxel-level threshold, and a p-FWE < 0.05 cluster size threshold^77^, as recommended^78,79^ and commonly applied^80–82^. A left ANT seed, a mirror location in the right ANT, and a parvocellular division of left mediodorsal thalamic nucleus seed for the DMNT were used for seed-to-voxel functional connectivity analysis (Table 1). The MNI coordinates of the seeds were those reported in an fMRI study examining BOLD activity during a recognition and cued recall task using atlas-based regions of interest^30^. The locations chosen are described below.

**Table 1.** MNI coordinates for seed locations (coordinates adapted from findings by Pergola *et al.*, 2013)^30^.

| Seed | Location | Side | x | y | z |
| --- | --- | --- | --- | --- | --- |
| 1 | anterior nuclei of the thalamus | L | -8 | -6 | 3 |
| 2 | anterior nuclei of the thalamus | R | 8 | -6 | 3 |
| 3 | dorsomedial nucleus of the thalamus | L | -6 | -19.5 | 7.5 |

Seed 2 was mirrored to the right side from the coordinates for the left side (Seed 1) by inverting the x-coordinates. ANT: anterior nuclei of the thalamus; MDpc = parvocellular division of mediodorsal thalamic nuclei; L: left; R: right

We contrasted the 3BPM with both the 3B and the 2BPM conditions. A voxel threshold of 0.001 was again applied. Where trends were observed, we explored these with a 0.005 threshold, which is commonly applied in PPI analyses^83,84^. We additionally compared the 3B and 2B conditions to evaluate whether connectivity differences involved similar locations when only the cognitive load was increased, without involving prospective memory. Note that the behavioral findings in the 2-back condition were consistent with strategic monitoring for the prospective memory task. Any functional connectivity differences in the 2BPM compared with the 2B condition would be expected to be continuous throughout the block, and therefore not related to prospective memory item onset times. As with the seed-based connectivity analysis above, individual time series were weighted by a boxcar signal characterizing each condition convolved with an SPM canonical hemodynamic response function. The weights were defined over the 2 s following prospective memory and ongoing task onset times.

#### Statistical analysis

A two-way repeated measures ANOVA was applied to the functional connectivity with each seed, with the factors *Task Type* (with, without prospective memory) and *N-Back* (2-back, 3-back), at voxel threshold < 0.001 and cluster size threshold p < 0.05, as per above.

##### Direct comparisons

Based on the results from the seed-to-voxel whole brain functional connectivity analysis, confirmatory analyses with direct comparisons were performed. We assessed ANT laterality, whether increased cognitive load accounts for the greater functional connectivity when prospective memory is involved, and specificity of the ANT in contrast with another thalamic nucleus engaged in declarative memory, the DMNT. Evaluations were performed by comparing left ANT functional connectivity with the regions-of-interest (ROIs) identified in the seed-to-voxel whole brain analyses. In the 2-back task, the ROI identified was the left DLPFC, and in the 3-back tasks, the left and the right STG (see Results: Functional connectivity section). We extracted the mean signal from the significant clusters and calculated functional connectivity in the conditions (2BPM, 3BPM, 2B, 3B) between which the clusters were found. Additionally, we used the mean signal from regions from the AAL3 atlas^58^ that encompassed those clusters.

###### Lateralization

To test for the laterality of the ANT connectivity variation depending on prospective memory, repeated measures ANOVAs were applied, with the within-subject factors *Laterality* (left ANT–ROI, right ANT–ROI) and *Task Type* (with, without prospective memory).

###### Cognitive load

To assess whether the greater left ANT–ROI functional connectivity seen in conditions involving prospective memory simply reflected a greater cognitive load, we applied repeated measures ANOVAs with the between-subject factors *N-back* (2-back, 3-back) and *Task Type* (with, without prospective memory) to the left ANT–ROI functional connectivity. We also applied two-sided paired T-tests to compare the functional connectivity difference between the n-back task alone and with a prospective memory component and the functional connectivity difference between the 2- and 3-back tasks alone.

###### Specificity of ANT

To test for the specificity of the PM-dependent connectivity difference on the thalamic nuclei, ANT, repeated measures ANOVAs were applied, with the within-subject factors *Thalamic Nuclei* (left ANT–ROI, left DMNT–ROI) and *Task Type* (with, without prospective memory) for the 2- and 3-back tasks.

## Results

### Behavioral findings

#### Approach to task

More participants reported mostly guessing in the 3-than the 2-back conditions (Fig. 2). Paired T-tests showed participants reported greater confidence in their ability to perform the 2-(score mean = 3.1, SD 1.2) than 3-back task alone (M = 2.4, SD 1.1; T(9) = 2.69, p = 0.025). These findings were mirrored when the prospective memory task component was included (2-back with prospective memory: M = 3.1, SD 1.2; 3-back with prospective memory (M = 1.8, SD 0.92; T(9) = 3.07, p = 0.013). Adding the prospective memory task component did not result in a significant difference in reported ability to perform the task in either the 2- or 3-back conditions (p’s > 0.05).

**Figure 2.**
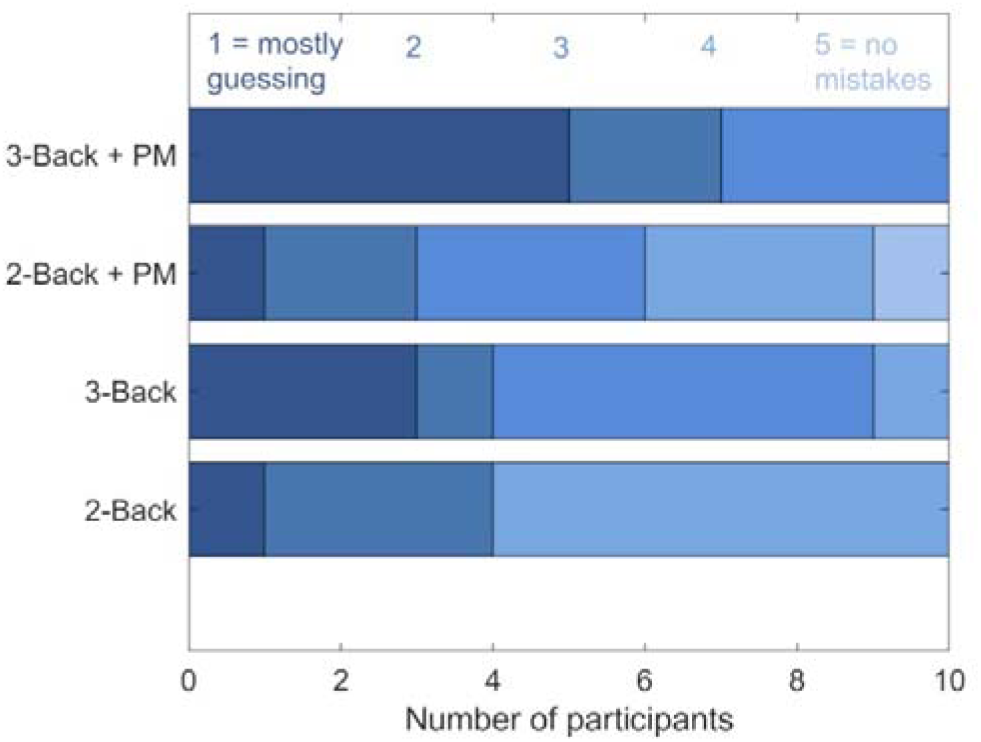
Participants’ ratings of their ability to perform the particular task type. PM: prospective memory component. Dark blue reflects score 1, progressing with the scale to light blue for score 5.

#### Task performance and cognitive load

Accuracy and RTs were normally distributed according to the Shapiro-Wilk test, with no outliers (defined as exceeding three times the interquartile range above the third or below the first quartile).

##### N-back task performance

Main effects of *N-Back* (F[1,13] = 30.4, p < 0.001, η_p_^2^ = 0.70) and *Task Type* (F[1,13] = 6.1, p = 0.029, η_p_^2^ = 0.32) were observed. Post hoc testing showed that d’ was greater during the ongoing task alone (mean d’ = 1.69, 95% CI = [1.47 1.91]) than when the prospective memory task was introduced (M = 1.56 [1.38 1.73]), and the d’ was greater during 2B (M = 1.87 [1.65 2.10] than 3B (M = 1.37 [1.17 1.58]). There was no significant interaction.

A main effect of *Task Type* (F[1,13] = 21.0, p < 0.001, η_p_^2^ = 0.62) was observed for RTs, but no main effect of *N-Back* and no interaction (p > 0.05). Post hoc testing showed that the RTs were faster during the ongoing task alone (mean RT = 946 ms, 95% CI = [880 1013]) than when the prospective memory task was introduced (mean RT = 1038 ms [974 1101]) (p < 0.001).

##### Prospective memory task component performance

Examining prospective memory responses, the percentage of responses to prospective memory items that were correct did not differ significantly between the 2-back (M = 61.7%, SD = 24.7) and 3-back (M = 61.2%, SD = 22.4) conditions (T(13) = 0.12, p = 0.90). The RTs also did not differ significantly between the 2-back (M = 926 ms, SD = 113) and 3-back (M = 874 ms, SD = 136) conditions (T(13) = −0.88, p = 0.39). Furthermore, the RTs did not differ significantly between ongoing task (M = 946 ms, SD = 115) and prospective memory (M = 900 ms, SD = 119) items (T(13) = 1.18, p = 0.26).

##### Effect of reported approach on N-back performance

When including *Reported Approach* as a covariate, a three-way interaction was observed between *Task Type*, *N-Back*, and *Reported Approach* (F(1,10) = 8.60, p = 0.015, η_p_^2^ = 0.46; Fig. 3). After correcting for *Reported Approach*, the two-way interaction remained between *Task Type* and *N-Back* (F(1,10) = 5.79, p = 0.037, η_p_^2^ = 0.37), and the effect was greater for participants reporting a strategic monitoring approach. Post hoc testing showed a lower d’ in the 2-back when the prospective memory was included (M = 1.80 [1.61 1.98]) than in the 2-back alone (M = 1.92, 95% CI = [1.68 2.16]; p = 0.036). No such difference was observed in the 3-back condition (3-back with prospective memory: M = 1.31 [1.03 1.59]; 3-back alone: M = 1.41 [1.23 1.59]; p = 0.16). The d’ was lower with than without the prospective memory task in the 2-back condition (p < 0.001) and in the 3-back condition (p = 0.003). Applying the analysis to RTs, no three-way interaction was seen, but an analogous tendency was observed examining a potential interaction between *Task Type* and *N-Back* after correcting for *Reported Approach* (F(1,10) = 2.59, p = 0.14, η_p_^2^ = 0.21). Post hoc testing was applied to examine whether the direction of any RT differences suggested a speed–accuracy trade-off, or supported the accuracy findings. RTs were slower in the 2-back when the prospective memory component was included (M = 948 ms [855 1041]) than in the 2-back alone (M = 904 ms [843 965]; p = 0.23). Again, like for accuracy, the difference was less in the 3-back condition (3-back with prospective memory: M = 1019 ms [927 1111]; 3-back alone: M = 1020 ms [961 1077]; p = 0.97). The RTs were slower with than without the prospective memory task in the 2-back condition (p < 0.001) and in the 3-back condition (p = 0.045).

**Figure 3.**
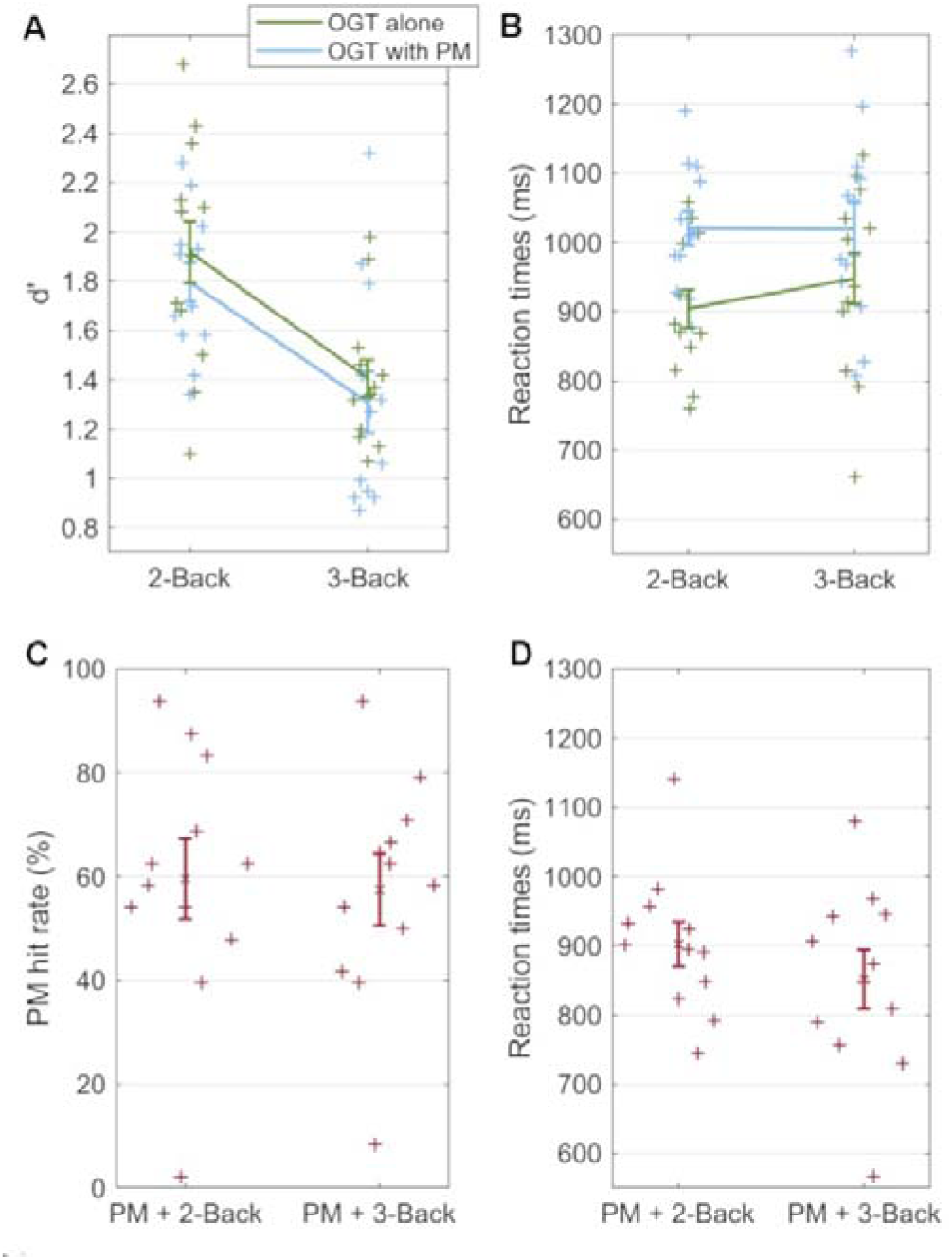
Behavioral performance during the task, reflected in accuracy (d’) and reaction times. Individual participant data points and standard error of the mean are shown. **A**. Accuracy in the n-back task as the ongoing task with different cognitive loads, with and without a prospective memory component. ANOVA: *Task Type* x *N-Back* interaction after correcting for *Reported Approach* (F(1,10) = 5.79, p = 0.037, η_p_^2^ = 0.37), with greater effects for participants reporting employing strategic monitoring. Post hoc: lower accuracy in the 2-back when the prospective memory was included than in the 2-back alone (p = 0.036). Accuracy was lower with than without the prospective memory task in the 2-back (p < 0.001) and in the 3-back condition (p = 0.003). **B**. Reaction times during the ongoing task with different cognitive loads, alone and with a prospective memory component included. ANOVA: no significant *Task Type* x *N-Back* interaction after correcting for *Reported Approach* (F(1,10) = 2.59, p = 0.14, η_p_^2^ = 0.21). **C**. Hit rate in the prospective memory task, with different cognitive loads in the ongoing task. ANOVA: *N-Back* x *Reported Approach* interaction (F(1,10) = 9.37, p = 0.012, η_p_^2^ = 0.48). Post hoc: faster RTs to prospective memory cues in the 3-back than the 2-back (p = 0.02) condition. **D**. Reaction times during the prospective memory task, with different cognitive loads in the ongoing task. ANOVA: no significant interactions or main effects. OGT: ongoing task; PM: prospective memory component.

##### Effect of reported approach on prospective memory performance

An interaction was seen in the RTs for prospective memory responses between *N-Back* and *Reported Approach* (F(1,10) = 9.37, p = 0.012, η_p_^2^ = 0.48; Fig. 3). Post hoc testing showed faster RTs to prospective memory cues in the 3-back (M = 851 ms, 95% CI = [766 938]) than the 2-back (M = 902 [835 970]; p = 0.02) condition. No interaction was observed for the prospective memory hit rate between *N-Back* and *Reported Approach* (F(1,10) = 0.84, p = 0.38, η_p_^2^ = 0.077).

##### Performance over time

Mean RT was faster at the end (M = 979 ms, SD = 1090) than the start of the blocks (M = 1034 ms, SD = 949; T(13) = 3.12, p = 0.008). Accuracy did not significantly differ between the start and end, and no significant interactions or main effects were observed for the factors *Task Type* and *N-Back* for the change in d’ or RT from the beginning to the end of the blocks (all p > 0.05).

### Functional connectivity

#### Seed 1: left ANT

Examining functional connectivity with Seed 1 in the ANT, clusters based on the applied significance thresholds (voxel threshold p < 0.001; cluster size threshold p < 0.05) were identified. The clusters reported were significant with a minimum T-value of T(13) = 4.22. Cluster locations where the cluster size p-FWE < 0.05 are given, together with cluster size, k.

A main effect of *Task Type* was observed in the left dorsolateral superior frontal gyrus according to the AAL3 atlas, in the DLPFC based on the cluster coordinates, and in the left superior temporal gyrus (STG) (Table 2). Based on the effect sizes, the functional connectivity with each cluster location was greater in the prospective memory condition than in the ongoing task alone (Fig. 1C, D).

**Table 2.** Functional connectivity with the left ANT (Seed 1) according to *Task Type* and *N-Back*.

| MNI-coordinates | Cluster location | Brodmann's area<sup>108</sup> | Cluster size | T(13) | p-FWE |
| --- | --- | --- | --- | --- | --- |
| Main effect of *Task Type* |  |  |  |  |  |
| -22 38 42 | left dorsolateral superior frontal gyrus (DLPFC) | BA 9 | 39 | 7.07 | 0.0070 |
| -54 -28 10 | left superior temporal gyrus | BA 22 | 33 | 7.65 | 0.019 |
| Main effect of *N-Back* |  |  |  |  |  |
| 06 44 00 | right anterior prefrontal cortex | BA 10/11 | 63 | 8.01 | 0.0018 |
| -16 -44 70 | left postcentral gyrus: primary somatosensory cortex | BA 2 | 42 | -7.58 | 0.0047 |
| Interaction between *Task Type* and *N-Back* |  |  |  |  |  |
| 52 24 22 | right inferior frontal gyrus | BA 45 | 33 | -6.87 | 0.021 |
| 24 -98 -08 | right inferior occipital gyrus | BA 18 | 32 | -5.74 | 0.025 |
| -50 -70 26 | left angular gyrus | BA 39 | 30 | -7.5 | 0.035 |
| 04 66 -18 | near right medial orbital superior frontal gyrus | BA 48 | 29 | -6.20 | 0.042 |
| Post hoc test of main effect of *Task Type*: 2BPM vs. 2B |  |  |  |  |  |
| -16 42 44 | left dorsolateral superior frontal gyrus (DLPFC) | BA 9 | 59 | 7.85 | 0.00028 |
| Post hoc test of main effect of *Task Type*: 3BPM vs. 3B: 3BPM > 3B |  |  |  |  |  |
| -48 -30 06 | left superior temporal gyrus | BA 48 | 143 | 10.26 | <0.000001 |
| 52 -18 04 | right superior temporal gyrus | BA 48 | 75 | 10.55 | 0.000037 |
| Post hoc test of main effect of *Task Type*: 3BPM vs. 3B: 3B > 3BPM |  |  |  |  |  |
| 4 40 50 | right medial superior frontal gyrus | BA 8 | 45 | -7.91 | 0.0030 |
| 48 24 20 | right inferior frontal gyrus | BA 48 | 44 | -10.10 | 0.0035 |
| 44 -78 22 | right middle occipital gyrus | BA 39 | 31 | -7.13 | 0.031 |
| Post hoc test of main effect of *Task Type*: 3B vs. 2B |  |  |  |  |  |
| 04 46 00 | right medial superior frontal gyrus | BA 10/11 | 50 | 6.91 | 0.0012 |
| 50 10 50 | right middle frontal gyrus | BA 9/44 | 30 | 6.02 | 0.035 |
| PPI: 3BPM vs. 3B |  |  |  |  |  |
| -62 -36 26 | left supramarginal/superior temporal gyrus | BA 48 | 19 | 6.72 | 0.00038 |
| 66 -24 26 | right supramarginal/superior temporal gyrus | BA 2/48 | 9 | 5.53 | 0.092 |
| PPI: 3BPM vs. 3B, with voxel threshold $p = 0.005$ | | | | | |
| -62 -36 26 | left supramarginal/superior temporal gyrus | BA 48 | 53 | 6.49 | 0.000034 |
| 66 -24 26 | right supramarginal/superior temporal gyrus | BA 2/48 | 21 | 5.38 | 0.099 |
| PPI: 3BPM vs. 2BPM |  |  |  |  |  |
| -60 -36 24 | left supramarginal/superior temporal gyrus | BA 48 | 9 | 8.28 | 0.12 |
| PPI: 3BPM vs. 2BPM, with voxel threshold $p = 0.005$ | | | | | |
| -60 -36 24 | left supramarginal/superior temporal gyrus | BA 48 | 47 | 8.00 | 0.00019 |
| -26 +66 48 | right frontal area (not an AAL3 region) | not a BA region | 21 | -6.91 | 0.012 |
| PPI: 3B vs. 2B, with voxel threshold $p = 0.005$ | | | | | |
| 26 -96 -10 | right inferior occipital gyrus | BA 18 | 32 | -6.51 | 0.0080 |

PPI (psychophysiological interaction) refers to the additional analysis in which trial onset was included. Voxel threshold $p = 0.001$ , unless otherwise specified. 2B: 2-back task; 2BPM: 2-back task
with prospective memory component; 3B: 3-back task; 3BPM: 3-back task with prospective memory component; BA: Brodmann's area

While a main effect of *N-Back* was also observed, the cluster locations differed from those showing a main effect of *Task Type*. Clusters were located in the right anterior prefrontal cortex and the left postcentral gyrus or somatosensory association cortex (Table 2).

An interaction between *Task Type* and *N-Back* was observed at the right inferior frontal gyrus, the right inferior occipital cortex, the left angular gyrus, and near to the right medial orbital superior frontal gyrus (Table 2).

Based on the interaction, post hoc comparisons were made. Contrasting the 2BPM with the 2B condition, functional connectivity was greater with left DLPFC, in a similar location to that identified as showing a main effect of *Task Type* (Fig. 4).

**Figure 4.**
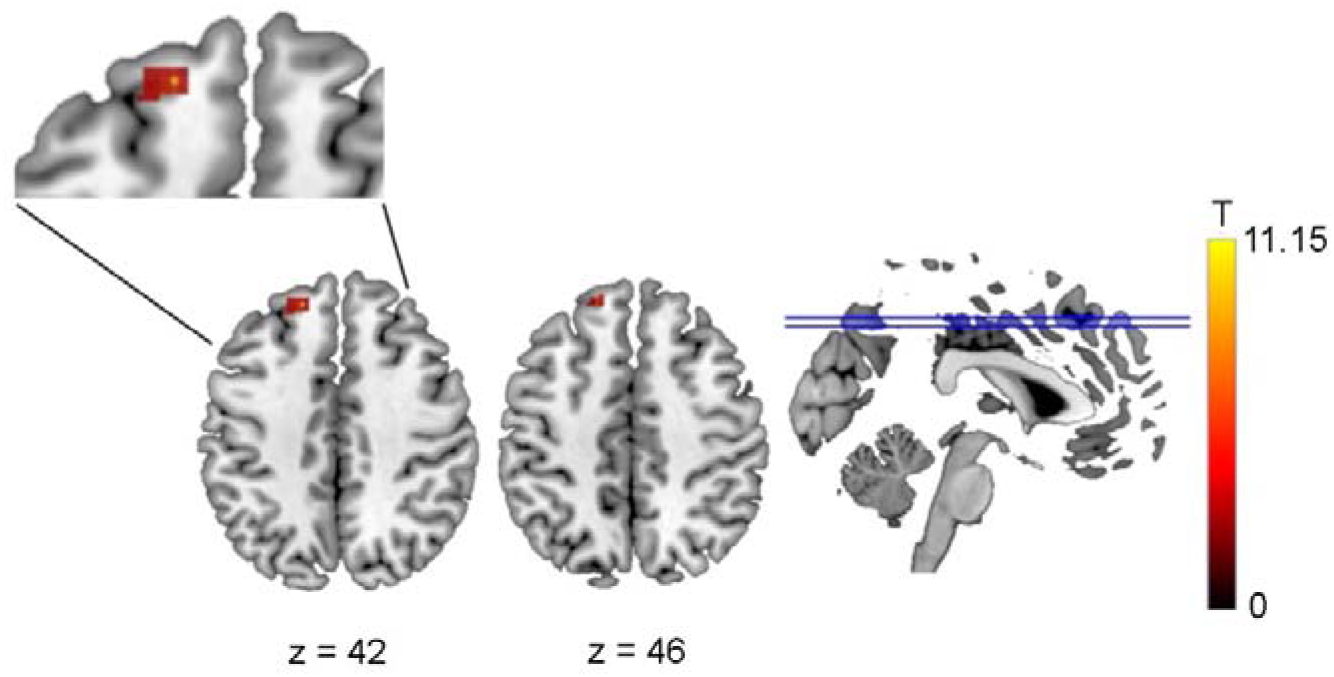
Functional connectivity difference between 2BPM and 2B conditions based on a seed in the anterior nuclei of the thalamus (Seed 1) Greater functional connectivity was observed in the 2BPM than the 2B condition with the left dorsolateral superior frontal gyrus (ANOVA: p-FWE = 0.00028). 2B: 2-back task; 2BPM: 2-back task with prospective memory component. T represents the T-values relating to the cluster size threshold.

Contrasting the 3BPM with the 3B condition, greater functional connectivity was observed in the 3BPM than the 3B condition in the left and right STG, in a similar location to that identified as showing a main effect of *Task Type* (Fig. 5). Greater functional connectivity in the 3B than 3BPM condition was seen with the right medial superior frontal gyrus, the inferior frontal gyrus, and the right middle occipital gyrus.

**Figure 5.**
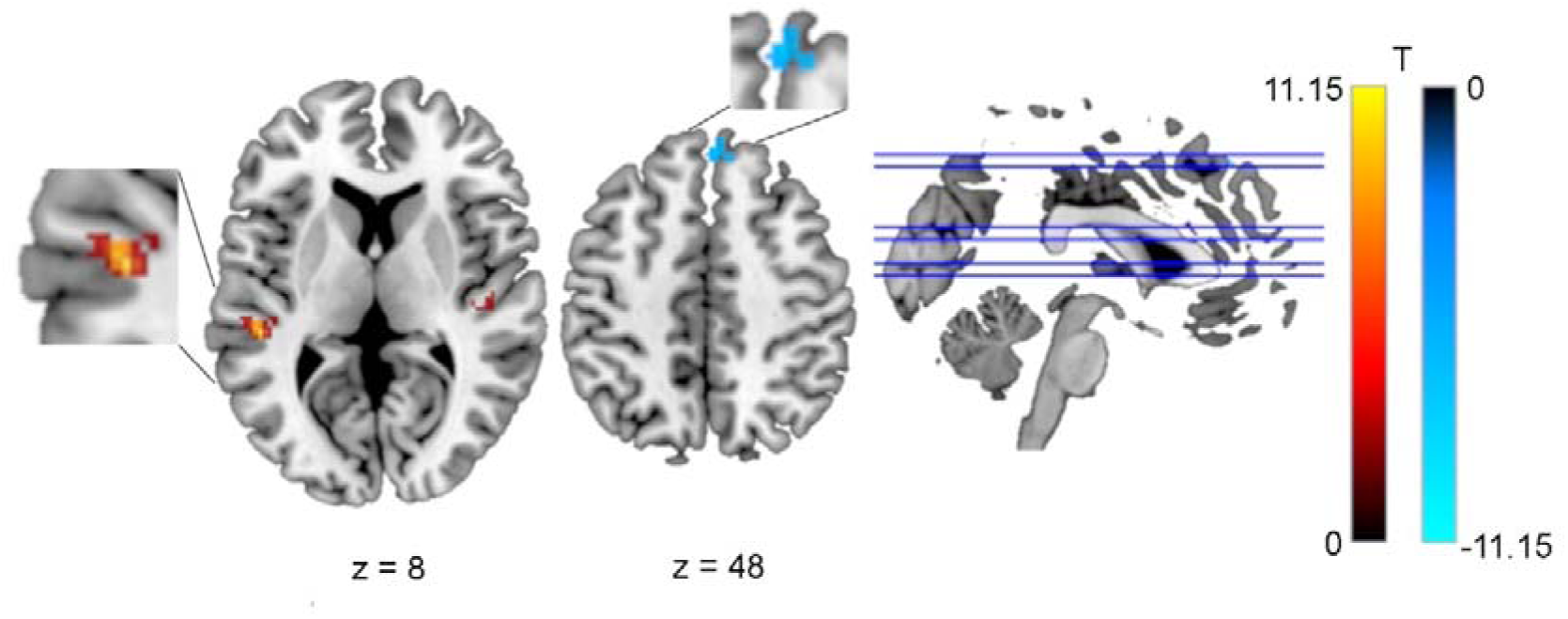
Functional connectivity difference between 3BPM and 3B conditions based on a seed in the anterior nuclei of the thalamus (Seed 1) Greater functional connectivity was observed in the 3BPM than the 3B condition in the in the left (p-FWE < 0.000001) and right superior temporal gyri (ANOVA: p-FWE = 0.000037). Greater functional connectivity in the 3B than 3BPM condition was seen with the right medial superior frontal gyrus (ANOVA: p-FWE = 0.0030), the inferior frontal gyrus (ANOVA: p-FWE = 0.0035), and the right middle occipital gyrus (p ANOVA: -FWE = 0.031). 3B: 3-back task; 3BPM: 3-back task with prospective memory component. T represents the T-values relating to the cluster size threshold.

Contrasting the 3B and 2B conditions, greater functional connectivity was seen in the 3B than the 2B condition with the right medial superior frontal gyrus and the right middle frontal gyrus (Fig. 6).

**Figure 6.**
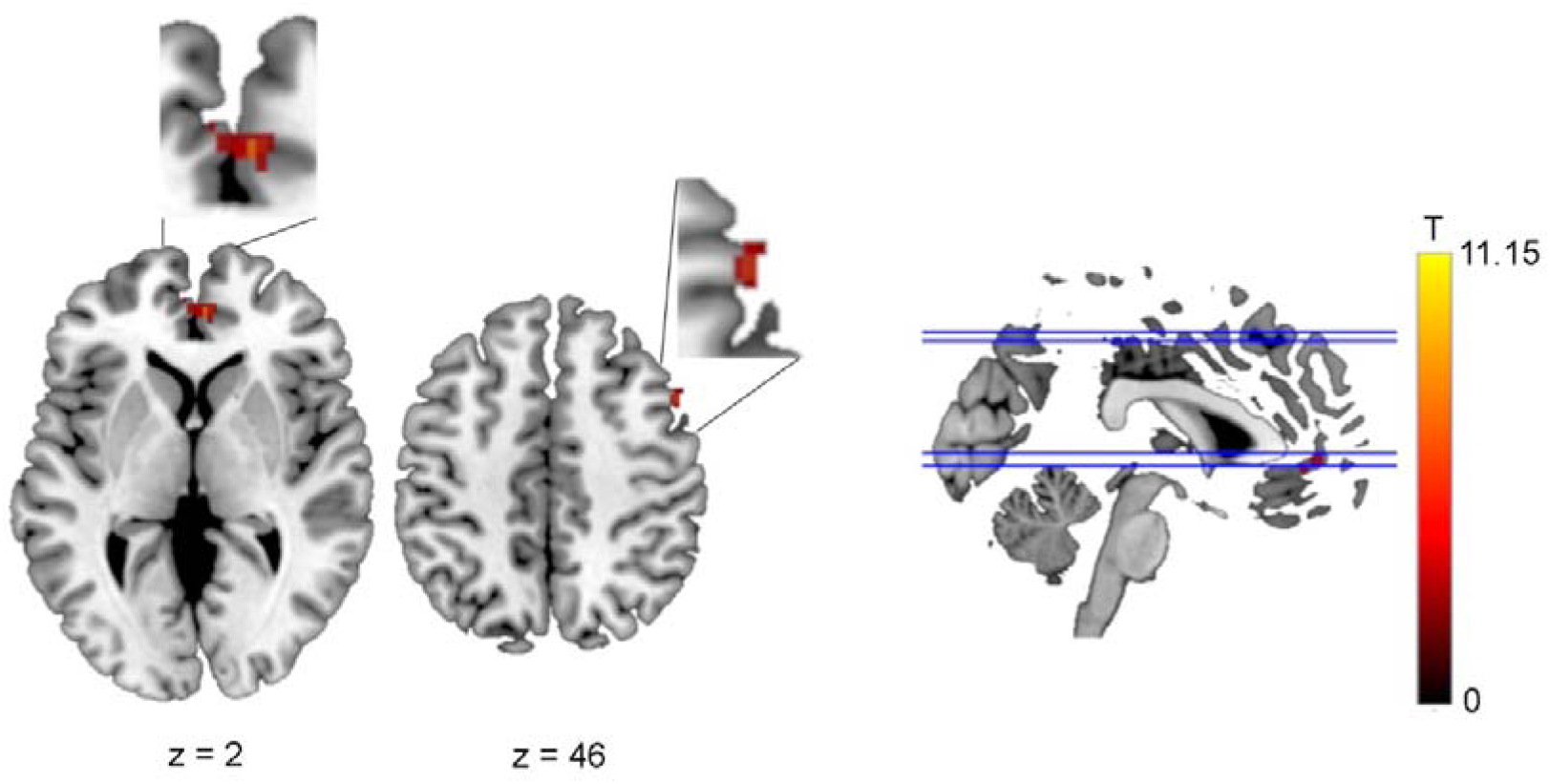
Functional connectivity difference between 3B and 2B conditions. Greater functional connectivity was observed in the right medial superior frontal gyrus (ANOVA: p-FWE = 0.0012) and in the right middle frontal gyrus (ANOVA: p-FWE = 0.035). 2B: 2-back task; 3B: 3-back task. T represents the T-values relating to the cluster size threshold.

Finally, contrasting 3BPM with 2BPM, no cluster was significant following FWE correction.

We note that while post hoc testing showed a significant difference between 2B and 2BPM and also between 3B and 3BPM, the effect size was smaller in the latter contrast.

The regression analysis including trial onset times showed greater functional connectivity during the 3BPM compared with the 3B condition with the left supramarginal/superior temporal gyrus and a trend to being greater with the right supramarginal/superior temporal gyrus. At the lower voxel threshold, the significance of the left temporal cluster increased, but that on the right remained a trend. 3BPM compared with 2BPM showed a trend towards greater connectivity with the left supramarginal/superior temporal gyrus only at a voxel threshold of 0.001 and was significantly greater at a voxel threshold of 0.005. Contrasting 3B and 2B conditions, no difference in connectivity with a temporal cortical cluster was observed.

#### Seed 2: right ANT

Applying an analogous ANOVA based on functional connectivity with the right ANT seed (Seed 2) also showed a main effect of *Task Type*, with clusters showing greater functional connectivity when a prospective memory component was included than when the ongoing task was performed alone, at a location near to the superior/middle/inferior temporal gyri on the same side, and less functional connectivity at the pallidum, near the left ANT (Table 3). A main effect of *N-Back* was also observed, with clusters in the right cuneus, the left calcarine fissure, the right inferior temporal gyrus, and the right cerebellum. An interaction was seen, with a cluster in the left dorsolateral superior frontal gyrus (Table 3).

**Table 3.** Functional connectivity with the right ANT (Seed 2) according to *Task type* and *N-Back*.

| MNI-coordinates | Cluster location | Brodmann's area | Cluster size | T(13) | p-FWE |
| --- | --- | --- | --- | --- | --- |
| Main effect of *Task Type*: with > without prospective memory |  |  |  |  |  |
| 48 -26 -10 | near right superior/middle/inferior temporal gyri | BA 48 | 37 | 9.16 | 0.0096 |
| Main effect of *Task Type*: without > with prospective memory |  |  |  |  |  |
| -12 6 -4 | left pallidum (near left ANT) | - | 83 | 7.36 | 0.00001 |
| Main effect of *N-Back* |  |  |  |  |  |
| 10 -88 14 | right cuneus | BA 18 | 86 | -12.1 | 0.000006 |
| -8 -90 -4 | left calcarine fissure | BA 17 | 64 | -8.11 | 0.00012 |
| 60 -38 -20 | right inferior temporal gyrus | BA 20 | 32 | 8.63 | 0.021 |
| 26 -58 -46 | right cerebellum | BA 48 | 30 | 8.64 | 0.031 |
| Interaction between *Task Type* and *N-Back* |  |  |  |  |  |
| -12 56 28 | left dorsolateral superior frontal gyrus | BA 9 | 36 | 6.33 | 0.0094 |
| Post hoc test of main effect of *Task Type*: 2BPM vs. 2B: 2BPM > 2B |  |  |  |  |  |
| -32 -28 40 | near left postcentral gyrus | - | 30 | -9.38 | 0.032 |
| Post hoc test of main effect of *Task Type*: 3B vs. 2B |  |  |  |  |  |
| -04 12 12 | near left caudate | - | 32 | 8.76 | 0.022 |
| -6 52 8 | left medial superior frontal gyrus | BA 10 | 28 | 6.78 | 0.047 |
| Post hoc test of main effect of *Task Type*: 3BPM vs. 3B |  |  |  |  |  |
| 22 -6 02 | right pallidum | BA 48 | 43 | 8.34 | 0.0031 |
| Post hoc test of main effect of *Task Type*: 3BPM vs. 2BPM |  |  |  |  |  |
| 12 -90 4 | right calcarine fissure | BA 17 | 32 | 7.56 | 0.018 |
| -48 26 24 | left inferior frontal gyrus | BA 48 | 29 | -8.2 | 0.032 |

2B: 2-back task; 2BPM: 2-back task with prospective memory component; 3B: 3-back task; 3BPM: 3-back task with prospective memory component; BA: Brodmann's area

Based on the interaction, post hoc pairwise comparisons were performed (Table 3). Contrasting 2BPM with 2B, a cluster was seen in the contralateral (left) postcentral gyrus, with lower functional connectivity with Seed 2 in the prospective memory condition. Contrasting the 3B and 2B conditions, greater functional connectivity was seen in the 3B condition in a region close to the left caudate and in the left medial superior frontal gyrus. Contrasting the 3BPM and 3B conditions, greater functional connectivity was seen in the 3BPM condition in a region close to the right pallidum. Finally, contrasting 3BPM with 2BPM, clusters were seen in the right calcarine fissure and the left inferior frontal gyrus.

#### Seed 3: left DMNT

An analogous ANOVA based on the left DMNT seed (Seed 3) revealed a main effect of *Task Type*, with clusters in the right inferior frontal gyrus, around the left pre- and post-central gyrus, and the left superior temporal gyrus, and a main effect of *N-Back*, with a cluster in the left middle occipital gyrus (Table 4). No significant interaction was seen.

**Table 4.** Functional connectivity with the left DMNT (Seed 3) according to *Task Type* and *N-Back*.

| MNI-coordinates | Cluster location | Brodmann's area | Cluster size | T(13) | p-FWE |
| --- | --- | --- | --- | --- | --- |
| Main effect of *Task Type*: with > without prospective memory |  |  |  |  |  |
| 58 34 08 | right inferior frontal gyrus | BA 45 | 45 | 7.98 | 0.0027 |
| -54 -12 46 | near left pre-/ post-central gyrus | BA 4 | 41 | 9.62 | 0.0052 |
| -44 0 -8 | left superior temporal gyrus | BA 48 | 29 | 8.98 | 0.042 |
| Main effect of *N-Back* |  |  |  |  |  |
| -50 -78 6 | left middle occipital gyrus | BA 19 | 56 | -8.32 | 0.0004 |

BA: Brodmann's area

#### Direct comparisons

##### Lateralization

A significant 2×2 interaction was observed for functional connectivity with the left DLPFC between *Laterality* (left ANT, right ANT) and *Task Type* (with, without prospective memory) in the 2-back condition (Table S1). Post hoc tests showed significantly greater left ANT–DLPFC connectivity with than without prospective memory and significantly greater left ANT–DLPFC than right ANT–DLPFC connectivity when including prospective memory. The findings were analogous for functional connectivity with the left and the right STG in the 3-back condition, though only marginally significant for the latter comparison for the right STG.

##### Cognitive load

A significant 2×2 interaction was observed for left ANT–DLPFC functional connectivity between *N-back* (2-back, 3-back) and *Task Type* (with, without prospective memory) (Table S2). Post hoc tests showed significantly greater left ANT–DLPFC functional connectivity with prospective memory in the 2-back but not the 3-back task. If cognitive load were the sole driver of left ANT–DLPFC functional connectivity, analogous findings would be expected in the 3-back condition. On direct comparison, moreover, the connectivity was not significantly greater with prospective memory in the 3-than 2-back condition. Main effects should be viewed with caution in the presence of a significant interaction, but we note a significant main effect of *Task Type* (with, without prospective memory) but not of *N-back* (2-back, 3-back). Finally, the effect of an additional prospective memory task on top of the ongoing n-back task (i.e., 2-back with prospective memory > 2-back without) was relatively larger than the working memory load per se (i.e., 3-back > 2-back). The findings were analogous for functional connectivity between the left ANT and the left and the right STG, though for the left STG, when prospective memory was included, connectivity just reached significance for being greater in the 3-than 2-back condition.

##### Specificity of ANT

A significant 2×2 interaction was observed for ANT–DLPFC functional connectivity between *Thalamic Nuclei* (left ANT–DLPFC, left DMNT–DLPFC) and *Task Type* (with, without prospective memory) in the 2-back condition (Table S3). Post hoc testing showed significantly greater left ANT–DLPFC connectivity with than without prospective memory, which was not the case for left DMNT–DLPFC connectivity. Left ANT–DLPFC was greater than left DMNT–DLPFC connectivity with prospective memory. Analogous interactions were observed for connectivity with the left and the right STG, and connectivity was significantly greater with than without prospective memory. However, while left ANT–STG was greater than left DMNT–DLPFC connectivity with prospective memory, it was not significant for either the left or the right STG. In summary, the effect of prospective memory on thalamus connectivity was specific to the ANT.

##### Functional connectivity comparisons based on atlas-based ROI masks

These direct comparisons were repeated using atlas-derived ROIs encompassing the larger number of voxels defined as the DLPFC and STG regions (see Supplementary Material). Examining lateralization, the same interactions were observed (Table S4). Post hoc testing revealed analogous tendencies, which reached greatest significance for connectivity with the left STG. Similarly, in examining the influence of cognitive load, the functional connectivity of the left ANT was greater with all three cortical regions when prospective memory was added rather than a simple cognitive load increase from 2- to 3-back (Table S5). While the finding was a trend for the DLPFC, it was significant for the STG. Finally, the analogous interactions examining the left ANT compared with the left DMNT showed significant interactions for the STG (Table S6). For the DLPFC, there were significant main effects of which thalamic nucleus was examined and of whether prospective memory was added.

## Discussion

Supporting the hypothesis that the ANT play a role in coordinating prospective memory processing, the left ANT showed greater functional connectivity with the left DLPFC and STG when a prospective memory component was added to the ongoing working memory task. Functional connectivity between the left ANT and these regions was not observed when contrasting performance of the 2-back with the 3-back task, supporting the notion that the greater functional connectivity on introducing the prospective memory task does not simply reflect greater cognitive load. Direct comparisons underlined this finding, showing greater functional connectivity differences between the left ANT and these cortical regions when a prospective memory task was added to the 2-back task than when the working memory cognitive load was increased by performing a 3-back instead of 2-back task. This functional connectivity difference was not observed with the left DMNT, another thalamic nucleus engaged in declarative memory, pointing to a specific role for the ANT in prospective memory processing. Significant interactions were seen between the *Thalamic Nuclei* (left ANT–ROI vs. left DMNT–ROI) and whether the task included prospective memory, with significant post hoc differences on including prospective memory for connectivity with the left ANT but not the left DMNT. Finally, such functional connectivity was not observed for the right ANT, with direct comparison showing greater functional connectivity for the left than right ANT, when prospective memory was included, consistent with a left laterality in prospective memory.

### Task performance strategy

The deterioration in performance of the ongoing task, reflected in lower accuracy and slower RTs in the ongoing task on addition of the prospective memory component, suggests strategic monitoring for the prospective memory cue. Intentional attention allocation to the prospective memory task, when a strategic monitoring approach is taken, can interfere with ongoing task performance^12,23,85^. A non-focal task encourages strategic monitoring, reflected in greater cognitive resource control when coordinating processes at retrieval^15,86–88^. The non-focal nature of our task, in which the cognitive processing required to respond to a prospective memory cue differs from the working memory processes engaged in the ongoing task, is consistent with a strategic monitoring approach. The lower ongoing task accuracy during the 3-back than the 2-back condition, with no change in RTs, is consistent with increased cognitive load during the 3-back condition^89^. The decline in participants’ self-reported ability to perform the 3-compared with the 2-back tasks also fits with an increase in cognitive load. The correct response rate and RTs to prospective memory cues did not differ according to the ongoing task cognitive load, and moreover, participants did not report an impact on their ability to perform the task when a prospective memory component was added. Sustaining prospective memory performance in the 3-back condition, despite the negative impact on n-back performance and positively perceived ability to perform the task despite greater cognitive load, suggests that the approach used in achieving the prospective memory component in the 3-back condition differed to that employed in the 2-back condition, consistent with strategic monitoring for a prospective memory cue with the lower cognitive load and spontaneous retrieval when the cognitive load of the ongoing task was greater.

While we did not observe an impact of cognitive load (2-back < 3-back) on prospective memory performance, if such a difference existed, our power to detect it was within the range of 5 to 30%. The questionnaires were therefore an important element in our interpretations regarding the approach participants took to prospective remembering. Participants were asked whether they tended to rely more on spontaneous retrieval or strategic monitoring to perform the prospective memory task, and their responses were included as a covariate in the evaluation of accuracy (as d’ for the ongoing n-back task and as hit rate for the prospective memory task) and RT. Reporting of applying more strategic monitoring was associated with a greater negative impact on ongoing task performance in the low cognitive load 2-back condition. Reporting of more spontaneous retrieval was associated with a better performance in responding to the prospective memory task in the high cognitive load 3-back condition. On the assumption that a participant’s predominant approach resulted in better performance under the conditions most suited to that approach, these findings provide further support for the supposition that the low cognitive load condition enabled participants to maintain the prospective memory component of the task actively, through strategic monitoring, while a greater reliance on a spontaneous retrieval approach to the prospective memory task was more successful when the ongoing task had a high cognitive load. While this analysis provides support for the notion that strategic monitoring is favored in the low and spontaneous retrieval in the high cognitive load conditions, we nonetheless assume that individuals varied their approach according to the condition. Separate ratings for each condition would ideally involve a between-group study design, requiring a larger participant number, with different groups for the low and high cognitive load conditions. Such a design would enable participants to evaluate their approach for a condition without increasing the importance they placed on the prospective memory component in subsequent blocks. Emphasizing the importance of the ongoing task has been proposed as a key factor in enabling study of spontaneous retrieval^11^.

A further consideration is the use of the same four cue colors in all task types, which meant that after a color was assigned as a prospective memory cue, in subsequent blocks, it required an n-back response. We assumed that the emphasis on the n-back performance, as well as the cognitive load of the n-back task, would mean that the prospective memory instruction from a previous block would not be relevant in the next block. It has been suggested, however, that involuntary recall of a previous prospective memory instruction could result in a requirement of suppression of the prospective memory response on seeing that cue again^90,91^. Using a larger selection of colors, so that a color previously used as a prospective memory cue was not used in subsequent blocks would avoid this issue. However, using a unique color for prospective remembering would have made this cue more salient, reducing dependency on prospective memory that a different action was required for such a cue. Previous behavioral and imaging studies comparing post-prospective memory conditions with conditions in which no prospective memory component had been previously involved showed no differences, consistent with successful suppression of a previous prospective memory instruction^90,91^. Indeed, in everyday life example might be a petrol station serving as a prospective memory cue after forming a prospective intention to buy petrol, which importantly ceases to induce prospective memory retrieval after petrol has been bought. Commission errors, or failure to suppress prospective retrieval after the cue is no longer relevant, increase with age^90^. Such errors have been associated with prefrontal cortex activations interpreted as reflecting ongoing strategic monitoring^91^. Our observation of functional connectivity involving prefrontal cortex in the condition in which we assumed strategic monitoring took place, which was not observed in the condition in which spontaneous retrieval was thought to occur, is consistent with previous reports of successful suppression of previous prospective memory instructions in our young participant group.

The DLPFC has an established role in prospective memory processing^26^. Activation of region BA 9 in particular has been associated with prospective memory maintenance^10,92^, and this area has also previously been associated with prospective memory retrieval with a 2-back task as the ongoing task ^93^, as applied in the current study. Our finding of greater functional connectivity between the left ANT and BA 9 during prospective memory maintenance is therefore consistent with engagement of the ANT in coordinating the memory and attentional brain networks underpinning prospective memory. Greater functional connectivity was also observed in the condition involving prospective memory between the left ANT and the left STG as well as between the right ANT and right STG. The STG has been associated with top-down attentional processing and memory retrieval, and changes in functional connectivity between the STG and widespread cortical regions during stimulus maintenance in working memory have led to the proposal of the STG as a hub region^11,94,95^. Notably the right STG was implicated in the latter working memory study, which was suggested to be associated with the spatial nature of the particular working memory task used^95^. Modulation of STG activity has also been reported during processing of relationships between stimuli presented simultaneously^96^. This notion is interesting in the context of a prospective memory paradigm, in which a single stimulus is presented but is simultaneously relevant to two different tasks, the ongoing and the prospective memory tasks. Taken together, these findings are consistent with strategic monitoring for a prospective memory cue in the prospective memory condition, with engagement of top-down attention and retrieval of the prospectively encoded intention.

Examining post hoc comparisons of functional connectivity with the left ANT seed, performed on the basis of the significant interaction, the contrast between functional connectivity in the 2BPM and 2B conditions corresponded with the main effect of *Task Type* observed in functional connectivity with DLPFC, while the contrast between functional connectivity in the 3BPM and 3B conditions showed greater correspondence with the main effect of *Task Type* seen in functional connectivity with the STG. These findings are consistent with a greater memory load in the 3-back conditions and more capacity for strategic monitoring when the overall cognitive load is lower, in the 2-back condition. While the behavioral findings indicate successful prospective memory processing in the 3BPM condition, the functional connectivity results suggest that the prospective memory task could be accomplished via a different mechanism, with a greater reliance on spontaneous retrieval. This interpretation fits with reports of more engagement of temporal regions in spontaneous retrieval^11,91^. Moreover, a higher cognitive load can facilitate reliance on spontaneous retrieval, even in non-focal tasks, such as used in the current study, where strategic monitoring is usually involved^27,85,88^. We note that the effect size difference was greater contrasting the prospective memory with the ongoing task conditions using the 2-than the 3-back ongoing task. Given that the cluster sizes were greater and corresponding p-values smaller in the latter contrast, the greater overall effect size difference using the 2-back task is presumably due to the effect sizes reflecting a seed to whole brain functional connectivity analysis, with other brain regions contributing to the effect size difference without being included in any cluster exceeding the significance threshold.

### Lateralization

Greater functional connectivity between the left ANT and both the DLPFC and STG is consistent with a role for the left ANT in the brain networks supporting prospective memory. Direct comparison showed significant interactions between laterality and whether a prospective memory component was included in the task, with greater left than right ANT functional connectivity with all three cortical regions. The difference was a trend for the right STG, which may be because there is no direct anatomical connectivity between the left ANT and right STG.

The finding of laterality is in keeping with previous reports of left lateralization of activity related to prospective remembering, which was suggested to reflect the lateralization of language processing to the left hemisphere^27^. It also fits with the memory-related activation in the left ANT in the study from which we derived the seed^30^. Using the seed location mirrored to the right ANT showed a remarkably similar functional connectivity difference, however, with an ipsilateral (right) temporal cluster showing greater functional connectivity with the right ANT seed in the prospective memory condition than when the ongoing task was performed alone. We note that ANT–DLPFC functional connectivity was not observed with a right ANT seed, suggesting a laterality that could be attributed to the usage of letters in the n-back task and the lateralization of language in the left hemisphere in most right-handed individuals. We have previously observed modulation of electrophysiological activity in both the left and the right ANT during memory formation in a paradigm using visual scenes^36,38^. Given reports of laterality of ANT function and differences in functional connectivity here, engagement of left and right ANT in different functional networks is plausible, likely related to the modality of the paradigm, in which verbalization of the task is a potential strategy. Both ANT seeds showing greater functional connectivity with the STG does suggest engagement of the ANT on both sides in prospective remembering, however, albeit potentially in different ways.

### Cognitive load

Given that adding a prospective memory component has the potential to increase the cognitive load in the task, we also examined whether simply increasing the cognitive load of the ongoing task, by using a 3-rather than 2-back paradigm, would yield similar greater functional connectivity in contrast with the 2-back alone to that observed on adding the prospective memory component. While greater frontal functional connectivity was observed, the frontal areas that were involved, BA 10/11 and BA 9/44, were more medial and anterior in location than the region of BA 9 (part of the DLPFC), where including a prospective memory component was associated with greater functional connectivity than when the ongoing task was performed alone. Of note also is that the functional connectivity with these regions was on the right side rather than the left, suggesting indirect functional connectivity as a part of a wider network. On direct comparison, an interaction was seen between *N-back* (2-back, 3-back) and *Task Type* (with, without prospective memory). Left ANT–DLPFC functional connectivity was greater in the 2-back condition when a prospective memory component was included. When prospective memory was included, there was no significant difference between the 3- and 2-back task conditions, which would have been expected if the greater functional connectivity only reflected cognitive load. While the significant interaction limits interpretability of main effects, we note that there was a main effect of *Task Type*, with greater functional connectivity when prospective memory was included, but not of *N-back*, consistent with differing associated functional connectivity. Moreover, a greater difference was observed between the 2-back conditions with and without a prospective memory component than between the 3-back and 2-back conditions without prospective memory. Taken together, these observations are consistent with the effects on ANT–DLPFC functional connectivity reflecting prospective remembering beyond simply increasing the cognitive load. The finding is consistent with the suggestion that medial PFC is more active during an ongoing task requiring working memory, with the lateral PFC underpinning delayed intentions^26^. BA 11 is also active in working memory and the encoding of new information^97^. Encoding the next stimuli in a 3-back task could potentially require greater involvement of this region. BA 44, which is considered as part of the ventrolateral PFC, particularly on the right side, has been shown to be engaged in working memory and motor response inhibition^98^. Given the lower number of targets than non-targets in the ongoing task, inhibition of a non-target response could be greater in the more challenging 3-than 2-back task. Importantly, varying the cognitive load of the ongoing task alone did not produce the same functional connectivity patterns as introducing the prospective memory task component, suggesting that differences in connectivity on adding prospective memory do not simply reflect greater cognitive load.

Including a prospective memory component in the 3-back condition had no impact on ongoing task performance, consistent with spontaneous retrieval of prospective memory items when the cognitive load of the ongoing task was higher. If the greater ANT–STG functional connectivity in the 3BPM than 3B condition reflects spontaneous prospective memory retrieval, as opposed to a higher general cognitive load, this connectivity difference would be expected also to be detected when prospective memory onsets are included in a regression analyses. Greater connectivity was still observed between the left ANT seed and the left STG in the 3BPM compared with the 3B condition and also compared with the 2BPM condition. Moreover, no such connectivity was observed comparing the 3B and 2B only conditions, providing further support for the notion that the functional connectivity between the left ANT seed and the temporal cortex reflects processes underlying spontaneous prospective memory retrieval rather than reflecting a general increase in cognitive load. Direct comparisons identified significant interactions between simple cognitive load (2-vs. 3-back) and whether a prospective memory component was included. Main effects of whether prospective memory was included were observed but not of whether 2- or 3-back conditions were performed. A greater difference was found between the 2-back condition with and without prospective memory than between the 3-back and 2-back conditions without prospective memory. Taken together, these findings are consistent with prospective remembering having an impact on functional connectivity between the ANT and temporal cortex beyond simple cognitive load increase.

### Specificity of ANT

Lesion studies point to engagement of the thalamus in prospective memory processing. To examine the specificity of a role for the ANT, we also examined seed-to-voxel whole brain connectivity with a seed located in another thalamic nucleus with an established role in episodic memory, the DMNT^30,99^. The DMNT projects to frontal cortex, and dissociations have been reported between the roles of the DMNT and ANT in memory processing, with the DMNT thought to play a predictive role in memory encoding and an executive role in memory retrieval^29,30,34,35,99^. While functional connectivity between the DMNT and left frontal areas differed according to whether a prospective memory component was included in the task, the areas were not those usually reported to be engaged in prospective memory and did not overlap with those showing functional connectivity with the ANT, suggesting that the ANT play a specific role in prospective remembering. Furthermore, direct comparisons revealed interactions between the particular thalamic nucleus and whether a prospective memory component was included in the task, with greater functional connectivity between the left ANT and frontal and temporal cortex when a prospective memory component was included in the task.

### Functional connectivity comparisons based on atlas-based ROI masks

Additional connectivity analyses with atlas-based cortical ROIs showed similar though weaker effects, which was unsurprising as the ROIs were much larger and did not only encompass functionally relevant cortical subregions. (See Supplementary Material: Supplementary Discussion for a detailed report.) Functional specialization of cortical subregions for specific tasks has been well-established through electrocorticography^100^.

### Limitations

We tested a young study population, in whom prospective memory impairment was not expected, to establish the potential engagement of the ANT in cognitive processes supporting prospective memory. However, prospective memory impairment more commonly affects older age groups, and future work is required to examine whether deficits in the functional connectivity that we observed underpin the prospective memory deficits seen in older patient groups and those with specific neurological conditions. Given the multiple neuronal networks engaged in prospective memory, it is likely that disruption of differing processes underlies prospective memory impairment in different clinical populations.

Performance in an n-back task improves over time, reflected in faster RT, especially at the beginning^45^. To minimize the potential for the performance during one cognitive load to be influenced by having performed the task with the other cognitive load first, participants performed a practice session of each of the four block types outside the scanner first, also providing the opportunity to ask any questions. Furthermore, the order was counterbalanced over participants. We compared the change in accuracy and RT over the course of a given block type and found no main effect of cognitive load or of whether a prospective memory task component was included, suggesting that better performance with a low than high cognitive load and without than with the prospective memory cues reflects differences between the block types rather than training effects. Future work could avoid such a potential limitation, however, with a between-subject design, in which different participants were allocated to perform the different block types.

We aimed to exclude the possibility that the connectivity associated with the prospective memory conditions reflected a general increase in cognitive load by comparing with a contrast between two different cognitive loads without a prospective memory component and by including participants’ reported approach to task performance in the analyses. However, a replication of the findings with a different set of paradigms is required to evaluate whether this connectivity is consistently observed during prospective remembering, irrespective of the type of task.

A further consideration is the choice of task itself. We chose the n-back as the ongoing task, because multiple trials are performed and the timing of prospective memory retrieval is also known, both of which are important factors for studying functional connectivity using fMRI. However, showing the prospective memory cue multiple times within a relatively short time period increases the probability that the prospective memory task will be maintained in working memory, so that only strategic monitoring and not spontaneous retrieval is evaluated. While increasing the cognitive load of the ongoing task did mean that including the prospective memory component did not impair the ongoing task performance, suggesting that participants relied on spontaneous retrieval, and while some participants self-reported performing the prospective memory component through spontaneous retrieval, similar findings based on different paradigms are required to confirm our interpretation.

Finally, we consider the sample size as a potential limitation. While larger participant groups are commonly included in fMRI studies at lower magnetic field strengths, comparatively lower participant numbers are reported to be required in 7T-fMRI studies^47,101^. In studies directly comparing analyses of data recorded at different field strengths, the ultra-high spatial specificity at 7T enables examination of fMRI at an individual participant level, so that 7T studies can be performed with smaller participant numbers than when scanning at lower field strengths^102,103^. Moreover, a recent study comparing memory consolidation using 7T and 3T fMRI with nine participants per group revealed expected changes in functional connectivity at 7T field strength, which were not detectable at 3T, even when correcting for signal-to-noise ratio differences^104^. Differentiation between hippocampal subfields was possible in a group of fourteen participants during associative learning using 7T scanning^42^, and memory traces were detected in the medial temporal lobe in eight participants at 7T^41^. The contrast between 7T and 3T field strengths that showed better discernment, particularly of subcortical structures, at 7T, was performed with ten participants^40^. Differences in subcortical–cortical functional connectivity involving another small, subcortical structure, the subthalamic nucleus, and sensorimotor cortex were detected in a clinical study involving patient groups with twelve and 18 participants^105^. We note, however, that our a priori power analysis was based on a different study design to our own, as we were unable to identify a previous 7T study examining seed-based thalamic connectivity using a within-subject design. The reference study employed a between-rather than within-subject design. It may still be relevant in informing the sample size in our study, however, as within-subject study designs can, in principle, be more powered than between-subject designs^106^. For example, in fMRI studies, the influence of cardiac and respiratory factors is reduced^69^. However, this may not always be the case due to other potential factors, including order effect (which we minimized by pseudorandom order of conditions), correlated observations, and measurement precision. We note that while low power might limit the detection of additional functional connectivity differences, the observed effects in our study are robust, as evidenced by the statistically significant results at the chosen alpha level. Indeed, statistical significance in a smaller sample requires a stronger signal relative to noise^107^. Future work should investigate whether these findings are replicable with different prospective memory paradigms and larger sample sizes, to establish whether the ANT–cortical functional connectivity found here, when a prospective memory component was added to the ongoing task, reflects essential mechanisms underpinning prospective memory.

## Conclusion

Our findings support our hypothesis that the ANT have a coordinatory role in prospective memory processing. Increased functional connectivity was observed between the ANT and cortical areas previously reported to be engaged in prospective memory. Previous evidence points to an active role for the ANT in episodic memory and for the thalamus in general in prospective memory. Here we identified a specific co-activation of a frontal cortical area known to be associated with monitoring processes in prospective remembering and the ANT during prospective memory processing, casting light on the mechanisms underpinning the retrieval of prospectively encoded future intentions. The finding was not replicated in a contrast in which the cognitive load of the working memory task was increased without prospective memory, suggesting specificity for the complex combination of multiple systems underpinning prospective memory. Furthermore, the findings suggest that fMRI at an ultra-high field strength enables evaluation of functional connectivity with small, subcortical structures, including thalamic nuclei. Future studies, with larger participant groups, are nonetheless required to confirm these findings. A pivotal role for the ANT has potential implications both for patients with anterior thalamic lesions as well as for those receiving implantation of electrodes in the ANT for deep brain stimulation to treat pharmacoresistant epilepsy. ANT lesions can arise through Korsakoff’s syndrome following Wernicke’s encephalopathy as well as through specific anterior thalamic infarction^31^, and an impact of such conditions on prospective remembering has implications for subsequent rehabilitation programs. Moreover, a potential modulation of prospective remembering through ANT stimulation may require consideration when determining the optimal target stimulation site to maximize seizure reduction while minimizing effects on prospective memory.

## Data availability

The behavioral and imaging (DICOM format) data that support the findings of this study are available upon reasonable request.

## Acknowledgements

The authors would like to thank Dr. Claus Tempelmann for advice and assistance in design of the MRI data acquisition protocol.

