## Supplementary material for "Exploring anterior thalamus functional connectivity with cortical regions in prospective memory with ultra-high-field fMRI"

**Supplementary Results**

***Functional connectivity comparisons based on atlas-based ROI masks***

##### Lateralization

The direct comparisons were additionally performed using ROIs derived from the larger number of voxels defined as DLPFC in the AAL3 atlas^1^. The 2x2 interaction for functional connectivity with the left DLPFC between *Laterality* (left ANT, right ANT) and *Task Type* (with, without prospective memory) in the 2-back test did not reach significance (Table S4). There was a main effect of *Task Type* but not of *Laterality*. The DLPFC, as defined in the AAL3 atlas, includes a large cortical region, which has functionally specialized subregions with differing anatomical connectivity and electrophysiology^2–4^. BA 9, a part of the DLPFC, has been shown to be engaged in prospective memory^5^. We therefore also repeated the analyses of functional connectivity involving the DLPFC using the smaller DLPFC mask from the Brodmann atlas^6^, limited to BA 9. A 2x2 interaction was then observed between *Laterality* and *Task Type*. Analogously to our findings based on the cluster identified in the seed-to-voxel whole brain analysis, post hoc testing showed significantly greater left ANT–DLPFC connectivity with than without prospective memory and significantly greater left ANT–DLPFC than right ANT–DLPFC connectivity when including prospective memory. Returning to the AAL3 atlas to examine functional connectivity differences between left and right ANT and the left and the right STG, analogous significant interactions were observed to those found in the cluster-based analyses. Post hoc testing showed significantly greater left but not right ANT connectivity when prospective memory was included than without prospective memory, both with the left and also the right STG. Also, left ANT showed significantly greater connectivity than the right ANT with the STG, but the difference was only significant for the left STG.

##### Cognitive load

Using the AAL3 atlas, including prospective memory in the n-back task was associated with higher functional connectivity between the left ANT and the cortical ROIs than simply increasing the cognitive load by performing a 3- compared with 2-back task (Table S5). The difference was a trend for connectivity with the DLPFC and significant for connectivity with the left and the right STG.

##### Specificity of ANT

The 2x2 interaction between *Thalamic Nuclei* (left ANT, left DMNT) and *Task Type* (with, without prospective memory) in the 2-back test was not significant, when examining the whole DLPFC as defined in the AAL3 atlas (Table S6). There were significant main effects both of *Thalamic Nuclei* and *Task Type*. Examining analogous functional connectivity with the STG, significant 2x2 interactions were observed for both the left and the right STG. Post hoc testing showed greater functional connectivity with than without prospective memory, which was significant for the left STG.

**Supplementary Discussion**

*Functional connectivity comparisons based on atlas-based ROI masks*

With a view to informing future studies examining the role of the ANT in prospective memory, we also compared functional connectivity between the thalamic seeds and the cortical areas identified in the seed-to-voxel whole brain connectivity analysis, the DLPFC in the 2-back and the left and right STG in the 3-back task variant, using AAL3 atlas masks for the cortical areas. The *Thalamic Nuclei* x *Task Type* and *Laterality* x *Task Type* interactions that we identified using the seed-to-voxel whole brain connectivity analysis in the 2-back task variant were not significant for the left ANT–DLPFC functional connectivity. The DLPFC is not a uniform region, but rather exhibits functional specialization, with different regions showing different anatomical connectivity^2–4^. When using an atlas-based definition for the ROI, encompassing the entire DLPFC, functional connectivity associated with the task may have been masked by an absence of functional connectivity with voxels covering task-irrelevant regions of the DLPFC. Using the Brodmann atlas, a *Laterality* x *Task Type* interaction was observed, analogous to those based on the DLPFC-ROI derived from the seed-to-voxel whole brain connectivity analysis. The functional connectivity findings based on the AAL3 atlas definition of the STG were also consistent with those determined using the ROIs from the seed-to-voxel whole brain connectivity analyses. The same interactions were significant for the atlas-based ROIs. In the post hoc tests, connectivity was significantly greater for the left ANT but not the right ANT when prospective memory was included, except for the right STG, and directions were the same in the post hoc tests directly comparing the connectivity with the left and the right ANT. However, the post hoc tests were significant for the DLPFC and the left STG but not for the right STG.

The evaluation of whether the left ANT functional connectivity increases during prospective remembering simply reflected greater cognitive load also showed the same tendencies as those found in the seed-to-voxel whole brain connectivity analyses, reaching significance for the left ANT–left STG functional connectivity. Finally, the interactions between which thalamic nucleus was examined and whether prospective memory was included were significant for the left and right STG but not for the DLPFC. Again, the post hoc test directions were the same as those based on the seed-to-voxel whole brain analyses.

The ROIs are larger in the atlases than those identified in the seed-to-voxel whole brain analyses. Electrocorticography has provided evidence for functional specialization of cortical subregions for specific tasks^7^. If subregions of the DLPFC and STG interact with the ANT during prospective memory, including the entire atlas-based ROI would be expected to result in a smaller effect size. Future studies with further participants are required to evaluate this possibility.

**Supplementary Tables**

**Table S1.** Direct comparisons assessing ANT laterality in the functional connectivity identified between the left ANT and cortical clusters in the left DLPFC and the left and right STG. (ANT = anterior nuclei of the thalamus; DLPFC = dorsolateral prefrontal cortex, 2B = 2-back; 2BPM = 2-back with prospective memory component; 3B = 3-back; 3BPM = 3-back with prospective memory component; STG = superior temporal gyrus)

| ***Laterality* (left ANT–left DLPFC vs. right ANT–left DLPFC) × *Task Type* (2B vs. 2BPM)** | | |
| --- | --- | --- |
| Interaction | F(1,13) = 19.81 | p = 0.00065 |
| Post hoc | | |
| left ANT–DLPFC: 2BPM > 2B | T(13) = 6.00 | p = 0.000044 |
| right ANT–DLPFC: 2BPM > 2B | T(13) = 1.35 | p = 0.20 |
| 2BPM: left ANT–DLPFC > right ANT–DLPFC | (T(13) = 3.47 | p = 0.0041 |
| 2B: right ANT–DLPFC > left ANT–DLPFC | T(13) = 2.18 | p = 0.048 |
| ***Laterality* (left ANT–left STG vs. right ANT–left STG) × *Task Type* (3B vs. 3BPM)** | | |
| Interaction | F(1,13) = 9.88 | p = 0.0078 |
| Post hoc | | |
| left ANT–left STG: 3BPM > 3B | T(13) = 12.89 | p = 0.0000000088 |
| right ANT–left STG: 3BPM > 3B | T(13) = 2.69 | p = 0.018 |
| 3BPM: left ANT–left STG > right ANT–left STG | T(13) = 2.60 | p = 0.022 |
| 3B: right ANT–left STG > left ANT–left STG | T(13) = 1.85 | p = 0.088 |
| ***Laterality* (left ANT–right STG vs. right ANT–right STG) × *Task Type* (3B vs. 3BPM)** | | |
| Interaction | F(1,13) = 10.31 | p = 0.0068 |
| Post hoc | | |
| left ANT–right STG: 3BPM > 3B | T(13) = 6.77 | p = 0.000013 |
| right ANT–right STG: 3BPM > 3B | T(13) = 1.67 | p = 0.12 |
| 3BPM: left ANT–right STG > right ANT–right STG | T(13) = 1.93 | p = 0.075 |
| 3B: right ANT–right STG > left ANT–right STG | T(13) = 2.25 | p = 0.043 |

**Table S2.** Direct comparisons assessing whether increased cognitive load explains the greater functional connectivity identified between the left ANT and cortical clusters in the left DLPFC and the left and right STG when a prospective memory component was included in the n-back task. (ANT = anterior nuclei of the thalamus; DLPFC = dorsolateral prefrontal cortex, 2B = 2-back; 2BPM = 2-back with prospective memory component; STG = superior temporal gyrus)

| **Left ANT–DLPFC: *N-back* (2B vs. 3B) x *Task Type* (with vs. without prospective memory)** | | |
| --- | --- | --- |
| Interaction | F(1,13) = 17.23 | p = 0.0011 |
| Post hoc | | |
| left ANT–DLPFC: 2BPM > 2B | T(1,13) = 6.00 | p = 0.000044 |
| left ANT–DLPFC: 3BPM > 3B | T(1,13) = 0.88 | p = 0.40 |
| left ANT–DLPFC: 3B > 2B | T(1,13) = 4.20 | p = 0.0010 |
| left ANT–DLPFC: 3BPM > 2BPM | T(1,13) = 1.83 | p = 0.091 |
| Main effect: *Task Type* | | |
| with PM > without PM | T(1,13) = 4.18 | p = 0.0010 |
| Main effect: *N-back* | | |
| 3B > 2B | T(1,13) = 0.94 | p = 0.37 |
| (2BPM – 2B) > (3B – 2B) | T(1,13) = 2.38 | p = 0.033 |
| **Left ANT–left STG: *N-back* (2B vs. 3B) x *Task Type* (with vs. without prospective memory)** | | |
| Interaction | F(1,13) = 22.15 | p = 0.00041 |
| Post hoc | | |
| left ANT–left STG: 2BPM > 2B | T(1,13) = 0.12 | p = 0.90 |
| left ANT– left STG: 3BPM > 3B | T(1,13) = 12.89 | p = 0.0000000088 |
| left ANT– left STG: 2B > 3B | T(1,13) = 4.40 | p = 0.00072 |
| left ANT– left STG: 3BPM > 2BPM | T(1,13) = 2.23 | p = 0.044 |
| Main effect: *Task Type* | | |
| with PM > without PM | T(1,13) = 5.45 | p = 0.00011 |
| Main effect: *N-back* | | |
| 2B > 3B | T(1,13) = 0.44 | p = 0.67 |
| (3BPM – 3B) > (3B – 2B) | T(1,13) = 10.61 | p = 0.0000000090 |
| **Left ANT–right STG: *N-back* (2B vs. 3B) x *Task Type* (with vs. without prospective memory)** | | |
| Interaction | F(1,13) = 7.064 | p = 0.020 |
| Post hoc | | |
| left ANT–right STG: 2BPM > 2B | T(1,13) = 0.18 | p = 0.86 |
| left ANT– right STG: 3BPM > 3B | T(1,13) = 6.77 | p = 0.000013 |
| left ANT– right STG: 2B > 3B | T(1,13) = 2.66 | p = 0.020 |
| left ANT– right STG: 3BPM > 2BPM | T(1,13) = 1.42 | p = 0.18 |
| Main effect: *Task Type* | | |
| with PM > without PM | T(1,13) = 3.38 | p = 0.0049 |
| Main effect: *N-back* | | |
| 2B > 3B | T(1,13) = 0.88 | p = 0.40 |
| (3BPM – 3B) > (3B – 2B) | T(1,13) = 5.14 | p = 0.00019 |

**Table S3.** Direct comparisons assessing specificity of the ANT compared with another thalamic nucleus involved in declarative memory, the DMNT, in the functional connectivity identified between the left ANT and cortical clusters in the left DLPFC and the left and right STG. (ANT = anterior nuclei of the thalamus; DLPFC = dorsolateral prefrontal cortex, 2B = 2-back; 2BPM = 2-back with prospective memory component; 3B = 3-back; 3BPM = 3-back with prospective memory component; STG = superior temporal gyrus)

| ***Thalamic Nuclei* (****left ANT–DLPFC vs. left DMNT–DLPFC) × *Task Type* (2B vs. 2BPM)** | | |
| --- | --- | --- |
| Interaction | F(1,13) = 21.49 | p = 0.00047 |
| Post hoc | | |
| left ANT–DLPFC: 2BPM > 2B | T(13) = 6.00 | p = 0.000044 |
| left DMNT–DLPFC: 2BPM > 2B | T(13) = 1.21 | p = 0.25 |
| 2BPM: left ANT–DLPFC > left DMNT–DLPFC | T(13) = 5.29 | p = 0.00015 |
| 2B: left DMNT–DLPFC > left ANT–DLPFC | T(13) = 1.081 | p = 0.30 |
| ***Thalamic Nuclei* (left ANT–left STG vs. left DMNT–left STG) × *Task Type* (3B vs. 3BPM)** | | |
| Interaction | F(1,13) = 13.83 | p = 0.0026 |
| Post hoc | | |
| left ANT–left STG: 3BPM > 3B | T(13) = 12.89 | p = 0.0000000088 |
| left DMNT–left STG: 3BPM > 3B | T(13) = 1.96 | p = 0.071 |
| 3BPM: left ANT–left STG > left DMNT–left STG | T(13) = 1.18 | p = 0.26 |
| 3B: left DMNT–left STG > left ANT–left STG | T(13) = 2.70 | p = 0.018 |
| ***Thalamic Nuclei* (left ANT–right STG vs. left DMNT–right STG) × *Task Type* (3B vs. 3BPM)** | | |
| Interaction | F(1,13) = 14.72 | p = 0.0021 |
| Post hoc | | |
| left ANT–right STG: 3BPM > 3B | T(13) = 6.77 | p = 0.000013 |
| left DMNT–right STG: 3BPM > 3B | T(13) = 0.37 | p = 0.71 |
| 3BPM: left ANT–right STG > left DMNT–right STG | T(13) = 0.60 | p = 0.56 |
| 3B: left DMNT–right STG > left ANT–right STG | T(13) = 4.07 | p = 0.0013 |

**Table S4.** Direct comparisons assessing ANT laterality in the functional connectivity identified between the left ANT and the AAL3/Brodmann atlas regions in which cortical clusters were identified in the left DLPFC and the left and right STG. (ANT = anterior nuclei of the thalamus; DLPFC = dorsolateral prefrontal cortex, 2B = 2-back; 2BPM = 2-back with prospective memory component; 3B = 3-back; 3BPM = 3-back with prospective memory component; STG = superior temporal gyrus)

| ***Laterality* (left ANT–DLPFC vs. right ANT–DLPFC) × *Task Type* (2B vs. 2BPM): AAL3** | | |
| --- | --- | --- |
| Interaction | F(1,13) = 2.24 | p = 0.16 |
| Main effect: *Laterality* | | |
| left ANT–DLPFC > right ANT–DLPFC | T(13) = 1.68 | p = 0.14 |
| Main effect: *Task Type* | | |
| with PM > without PM | T(13) = 2.80 | p = 0.015 |
| ***Laterality* (left ANT–DLPFC vs right ANT–DLPFC) × *Task Type* (2B vs. 2BPM): Brodmann** | | |
| Interaction | F(1,13) = 4.71 | p = 0.049 |
| Post hoc | | |
| left ANT– DLPFC STG: 2BPM > 2B | T(1,13) = 2.90 | p = 0.012 |
| right ANT– DLPFC: 2B > 2BPM | T(1,13) = 0.016 | p = 0.99 |
| 2BPM: left ANT– DLPFC > right ANT– DLPFC | T(1,13) = 2.56 | p = 0.024 |
| 2B: right ANT– DLPFC > left ANT– DLPFC | T(1,13) = 0.28 | p = 0.79 |
| ***Laterality* (left ANT–left STG vs. right ANT–left STG) × *Task Type* (3B vs. 3BPM): AAL3** | | |
| Interaction | F(1,13) = 8.35 | p = 0.013 |
| Post hoc | | |
| left ANT–left STG: 3BPM > 3B | T(1,13) = 3.40 | p = 0.0048 |
| right ANT–left STG: 3BPM > 3B | T(1,13) = 0.66 | p = 0.52 |
| 3BPM: left ANT–left STG > right ANT–left STG | T(1,13) = 2.36 | p = 0.035 |
| 3B: right ANT–left STG > left ANT–left STG | T(1,13) = 1.30 | p = 0.20 |
| ***Laterality* (left ANT–right STG vs. right ANT–right STG) × *Task Type* (3B vs. 3BPM): AAL3** | | |
| Interaction | F(1,13) = 4.77 | p = 0.048 |
| Post hoc | | |
| left ANT–right STG: 3BPM > 3B | T(1,13) = 1.52 | p = 0.15 |
| right ANT–right STG: 3B > 3BPM | T(1,13) = 0.29 | p = 0.77 |
| 3BPM: left ANT–right STG > right ANT–right STG | T(1,13) = 1.14 | p = 0.28 |
| 3B: right ANT–right STG > left ANT–right STG | T(1,13) = 1.92 | p = 0.077 |

**Table S5.** Direct comparisons assessing whether increased cognitive load explains the greater functional connectivity identified between the left ANT and AAL3 atlas regions in which cortical clusters were identified in the left DLPFC and the left and right STG when a prospective memory component was included in the n-back task. (ANT = anterior nuclei of the thalamus; DLPFC = dorsolateral prefrontal cortex, 2B = 2-back; 2BPM = 2-back with prospective memory component; 3B = 3-back; 3BPM = 3-back with prospective memory component; STG = superior temporal gyrus)

| ***Cognitive Load*** | | |
| --- | --- | --- |
| left ANT–DLPFC: | | |
| (2BPM – 2B) > (3B – 2B) | T(1,13) = 1.54 | p = 0.15 |
| left ANT–left STG: | | |
| (3BPM – 3B) > (3B – 2B) | T(1,13) = 3.12 | p = 0.0081 |
| left ANT–right STG: | | |
| (3BPM – 3B) > (3B – 2B | T(1,13) = 2.12 | p = 0.054 |

**Table S6.** Direct comparisons assessing specificity of the ANT compared with another thalamic nucleus involved in declarative memory, the DMNT, in the functional connectivity identified between the left ANT and cortical clusters in the left DLPFC and the left and right STG, based on the regions in the AAL3 atlas in which the clusters were found. (ANT = anterior nuclei of the thalamus; DLPFC = dorsolateral prefrontal cortex, 2B = 2-back; 2BPM = 2-back with prospective memory component; 3B = 3-back; 3BPM = 3-back with prospective memory component; STG = superior temporal gyrus)

| ***Thalamic Nuclei* (left ANT–DLPFC vs. left DMNT–DLPFC) × *Task Type* (2B vs. 2BPM)** | | |
| --- | --- | --- |
| Interaction | (1,13) = 1.50 | p = 0.24 |
| Main effect: *Thalamic Nuclei* | | |
| left ANT–DLPFC > left DMNT–DLPFC | T(13) = 2.43 | p = 0.031 |
| Main effect: *Task type* | | |
| with PM > without PM | T(13) = 2.83 | p = 0.014 |
| ***Thalamic Nuclei* (left ANT–left STG vs. left DMNT–left STG) × *Task Type* (2B vs. 2BPM)** | | |
| Interaction | F(1,13) = 9.98 | p = 0.0075 |
| Post hoc | | |
| left ANT–left STG: 3BPM > 3B | T(1,13) = 3.40 | p = 0.0048 |
| left DMNT–left STG: 3BPM > 3B | T(1,13) = 0.02 | p = 0.98 |
| 3BPM: left ANT–left STG > left DMNT–left STG | T(1,13) = 0.66 | p = 0.52 |
| 3B: left DMNT–left STG > left ANT–left STG | T(1,13) = 3.00 | p = 0.0098 |
| ***Thalamic Nuclei* (left ANT–right STG vs. left DMNT–right STG) × *Task Type* (2B vs. 2BPM)** | | |
| Interaction | F(1,13) = 11.10 | p = 0.0054 |
| Post hoc | | |
| left ANT–right STG: 3BPM > 3B | T(1,13) = 1.52 | p = 0.15 |
| left DMNT–right STG: 3B > 3BPM | T(1,13) = 1.46 | p = 0.17 |
| 3BPM: left ANT–right STG > left DMNT–right STG | T(1,13) = 0.64 | p = 0.53 |
| 3B: left DMNT–right STG > left ANT–right STG | T(1,13) = 3.45 | p = 0.0043 |
